# Global patterns of species diversity and distribution in the biomedically and biotechnologically important fungal genus *Aspergillus*

**DOI:** 10.1101/2024.11.29.626055

**Authors:** Olivia L. Riedling, Kyle T. David, Antonis Rokas

## Abstract

*Aspergillus* fungi are key producers of pharmaceuticals, enzymes, and food products and exhibit diverse lifestyles, ranging from saprophytes to opportunistic pathogens. To improve understanding of *Aspergillus* species diversity, identify key environmental factors influencing their geographic distributions, and estimate the impact of future climate change, we trained a random forest machine learning classifier on 30,542 terrestrial occurrence records for 176 species (∼40% of known species in the genus) and 96 environmental variables. We found that regions with high species diversity are concentrated in temperate forests, which suggests that areas with mild seasonal variation may serve as diversity hotspots. Species range estimates revealed extensive variability, both within and across taxonomic sections; while some species are cosmopolitan, others have more restricted ranges. Furthermore, range overlap between species is generally low. The top predictors of mean species richness were the index of cumulative human impact and five bioclimatic factors, such as temperature and temperate vs non-temperate ecoregions. Our future climate analyses revealed considerable variation in species range estimates in response to changing climates; some species ranges are predicted to expand (e.g., the food spoilage and mycotoxin-producing *Aspergillus versicolor*), and others are predicted to contract or remain stable. Notably, the predicted range of the major pathogen *Aspergillus fumigatus* was predicted to decrease in response to climate change, whereas the range of the major pathogen *Aspergillus flavus* was predicted to increase and gradually decrease. Our findings reveal how both natural and human factors influence *Aspergillus* species ranges and highlight their ecological diversity, including the diversity of their responses to changing climates, which is of relevance to pathogen and mycotoxin risk assessment.

## Introduction

The genus *Aspergillus* is a highly diverse clade of filamentous fungi comprising around 450 described species distributed across 28 taxonomic sections (1,2). *Aspergillus* have been typically regarded as cosmopolitan since they have been isolated from various habitats across the globe—including soil, air, aquatic environments, Arctic regions, and living organisms (3). *Aspergillus* species have established their importance, particularly in the pharmaceutical, food science, agricultural, and cosmetic industries, thereby playing critical roles in human society (4). Examples of these bioeconomically important species include the industrial workhorse *Aspergillus terreus,* which produces the cholesterol-lowering pharmaceutical lovastatin (5), *Aspergillus niger*, a well-established cell factory for enzyme production (6), and *Aspergillus oryzae*, which is widely used in food manufacturing for fermented food products (7).

*Aspergillus* species occupy a wide array of ecological niches and exhibit diverse lifestyles. They can exist as saprophytes thriving on dead or decaying organic matter, endophytes, plant pests, and pathogens of humans and animals (8,9). Multiple species have been implicated in disorders in plants and plant products, but *Aspergillus niger* and *Aspergillus flavus* are the most common species identified in contaminated and spoiled agricultural products like fruits, vegetables, and nuts (10). Species in the genus also produce mycotoxins with adverse effects on health, like aflatoxins and ochratoxin A, both of which are known carcinogens (11,12). Some *Aspergillus* species are opportunistic pathogens that cause a range of diseases collectively known as aspergillosis (13). These infections can manifest in numerous locations, such as the lungs, skin, brain, and eyes (13,14), and affect over a million individuals globally each year. Invasive aspergillosis, one of the most severe forms of aspergillosis, results in very high mortality (14,15) and is primarily caused by the major pathogens *Aspergillus fumigatus* (15) and *A. flavus*, as well as approximately a dozen other minor pathogens (16). *Aspergillus* infections can also afflict a wide variety of animals, including mammals, birds, honey bees, fish, reptiles, and sea fan corals (17).

The diversity of lifestyles and ability of *Aspergillus* species to thrive in a range of environments suggests that species in the genus are ecologically highly diverse (18). A few reviews have summarized the isolation environments of *Aspergillus* species (4,9), but they focus on specific locations or regions with few species. In addition, global studies of fungal distributions have revealed climate-driven patterns, such as temperature and precipitation (19,20). However, these studies collapse the diversity and variation seen at lower taxonomic levels, such as genus-specific patterns in distributions and drivers of their distributions. Overall, species from the genus have seemingly been isolated across the globe in relatively stable to more extreme environments. Still, the general global patterns of the distributions of species genus-wide are largely unknown.

In recent years, numerous databases have been established to enhance fungal community sampling, such as the Global Biodiversity Information Facility (21) and the GlobalFungi database (22). Since comprehensive global sampling efforts are still in their infancy, one can utilize predictive algorithms to estimate species distributions. Traditionally, these algorithms have included generalized linear models and habitat suitability analyses, which use climate-related data—such as temperature and precipitation— and geographic features to predict distributions. These types of analyses have been primarily applied to estimate the distributions of species under changing climatic conditions. It is thought that climate change, particularly rising temperatures, could facilitate the expansion of fungal species ranges and ease the colonization of host organisms, thereby increasing the prevalence of emerging fungal pathogens (23–26). This topic has stimulated research into the distributions of various genera and species that impact human health or agriculture, including *Cryptococcus*, *Fusarium*, and *Coccidioides* (27–29).

In this study, we sought to examine the geographical distribution and diversity of species in the biomedically and biotechnologically important fungal genus *Aspergillus*. Specifically, we trained a random forest classifier (30) on 30,542 terrestrial occurrence records for 176 species (∼40% of known species in the genus) and 96 environmental variables to investigate the geographic distributions and ecological patterns of the genus *Aspergillus*. Additionally, we combined this framework with future climate models to predict the distributions of *Aspergillus* species under three climate change scenarios to explore differences between species distributions in response to changing climates. Our findings identify the diverse environmental factors that influence *Aspergillus* geographic ranges, such as natural and anthropogenic factors, and underscore the importance of continued monitoring of *Aspergillus* species to further both the understanding of the unique environmental dynamics impacting their ranges and to anticipate responses to the currently changing climates.

## Results and Discussion

### Areas of the highest species richness are centered in warm, humid, and temperate forests

To examine the distributions of geographic sampling of species obtained from the GlobalFungi database (22) across the genus, we mapped the sampling on a phylogeny of *Aspergillus*. The sampling spanned 27 of the 28 taxonomic sections (**Figure S1**), which enabled us to analyze global patterns of species diversity across the genus. Our data were from 17 different isolation sources, with soil being the largest source (soil: 61,724; topsoil: 23,748; rhizosphere soil: 6,029). To investigate the underlying patterns in species distributions, we used a random forest classifier to predict global species distributions for 236 *Aspergillus* species. We predicted distributions for species with > 4 occurrence records and excluded species with True Positive and True Negative rates < 75%, which yielded 176 species for further analyses (**Figure S2** and **Table S2**).

To identify areas with potentially high species diversity, we generated a species richness map (**Figure 1A**). We did not detect any latitudinal or longitudinal richness gradients across *Aspergillus* (**Figure S3**), consistent with data from other fungal lineages and with the raw data from GlobalFungi (**Figure S4**) (30–32). Areas of high richness include southeastern Europe, southeastern Asia, and portions of southwestern Africa, which are known for high biodiversity and more stable temperatures. Additionally, areas of moderate richness include portions of the United States, Europe, Central and South America, and Australia, which may have larger seasonal variations contributing to the reduced species richness. We compared the predicted hotspots of species richness to the sampling hotspots in the raw data and found that eastern Asia had a high sampling density and a high species richness. To address potential biases in sampling, we compared the relationship between sampling effort from the empirical observations in the training data with observed species richness and predicted species richness. We found the relationship between predicted richness and sampling effort (p = 8.24e-12, m = 0.24, r^2^ = 0.08) was much weaker than the observations in the training data (p = 9.13e-211, m = 0.84, r^2^ = 0.8) (**Figure S5**). Furthermore, there were several areas predicted to have moderate to high species richness from relatively low sampling, like portions of western Australia and South America.

**Figure 1:**
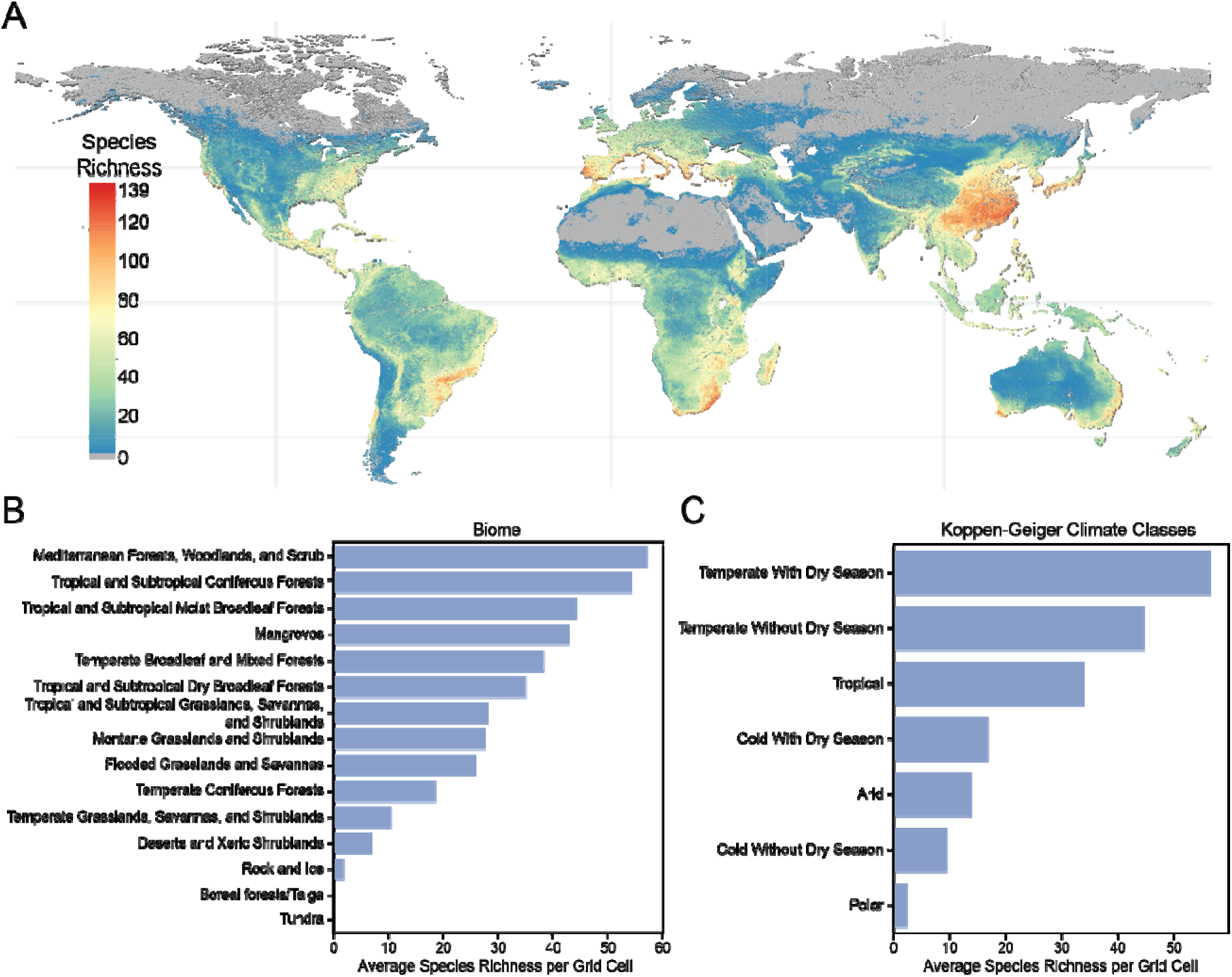
*Aspergillus* species richness is highest in temperate forests. A) Global map of *Aspergillus* species richness. Horizontal lines indicate latitudes of 40°N, 0°, and −40°S and vertical lines are longitudes of 100°W, 0°, and −100°E. Warmer colors display areas of higher species richness with a maximum of 139 species; colder colors are areas of lower species richness with a minimum of 1 species; and grey areas denote the absence of any species. B) Average species richness per biome. The X-axis is the average species richness per raster grid cell (pixel), and the Y-axis is the different biomes represented in the dataset ordered from highest to lowest average species richness. C) Average species richness per Köppen-Geiger (KG) climate classes, which classify climates based on temperature, precipitation, and seasonal patterns. The X-axis is the average species richness per raster grid cell, and the Y-axis is the different KG classes ordered from highest to lowest average species richness.

Species were distributed across 15 diverse biomes, with the highest average species richness (ASR) found in the Mediterranean Forests, Woodlands, and Scrub biome, which is characterized by its hot, dry summers and cool, moist winters and includes California, the Chilean Matorral, southern Africa, southwest and southern Australia, and the Mediterranean Basin (**Figure 1B**). Although this biome has the highest ASR, other biomes were also species-rich. For example, there were five other biomes that had an ASR of 30 species, illustrating the breadth of environments that *Aspergillus* species can occupy. We also analyzed the ASR across Köppen-Geiger climate classifications, which divide climates into major groups and subgroups according to seasonal precipitation and temperature (**Figure 1C**). We found that species richness is highest in temperate areas with dry seasons and temperate areas without dry seasons, specifically in the Temperate regions with Dry Winters and Warm Summers (**Figure S6**).

To identify specific environmental components associated with higher species richness, we analyzed ecofloristic zones (based on climate and vegetation type), soil classes, plant classes, and geomorphic classes (**Figure S7**). We found that the ecofloristic zones subtropical dry (ASR: 78) and humid (ASR: 77) forests, the soil class alisols (ASR: 77), the plant classes urban/built-up areas (ASR: 52) and deciduous broadleaf trees (average species richness: 38), and lastly the geomorphic class depression (ASR: 37) displayed the highest averages of species richness (**Figure S7**). The ecofloristic zone and soil class data further support the association of *Aspergillus* species richness with consistently warm, humid climate types (alisols are acidic and poorly drained soils typically found in humid tropical, humid subtropical, and humid temperate regions). Interestingly, our finding that the highest average richness in plant classes was found in urban/built-up areas reveals an association of *Aspergillus* with human-made environments. In summary, these results suggest that areas of higher *Aspergillus* species richness exhibit consistently warmer and more humid climates with little seasonal variation, have topographical variation, and contain human-made developments.

### Extensive variation in species ranges and overlap

To examine the size of species’ ranges and their overlap, we analyzed the percentage of grid cells or pixels of the raster image occupied by each species as a proxy for the extent of their geographic range. We found extensive variation in *Aspergillus* species ranges, with an average of 2.3% of global grid cells occupied per species (n = 176 species). *Aspergillus sigurros* (section *Usti*) and *Aspergillus flocculosus* (section *Circumdati*) had the largest predicted ranges of 4.83% and 4.51%, respectively, while *Aspergillus lucknowensis* (section *Usti*), *Aspergillus ambiguus* (section *Flavipedes*), and *Aspergillus unilateralis* (section *Fumigati*) had the noticeably smallest predicted ranges of 0.14%, 0.40%, and 0.52%, respectively (**Figure S8 and Table S4**). The ranges of *A. lucknowensis* and *A. ambiguus* are primarily restricted to the Mediterranean Forests, Woodlands, and Scrub biome. These results suggest that while some species have larger ranges and are cosmopolitan, others are more restricted and occupy specific environments.

To test whether the frequency of a species in the training data was associated with the size of their predicted ranges, we performed an ordinary least squares regression. We found a very weak, statistically significant negative relationship (p = 0.014, effect size = −0.0006, r² = 0.034) (**Figure S9A** and **Table S5**). Thus, while species with fewer occurrence records in the training data might have slightly larger predicted ranges, their overall impact on species range predictions is small compared to specific ecological and environmental factors.

To test whether species ranges differed by taxonomic section, we plotted the percentage of grid cells occupied for 163 / 176 species from 25 sections (with available nucleotide sequences for the three taxonomic maker genes β-tubulin, calmodulin, and RNA polymerase β) on the *Aspergillus* phylogeny. We found the ranges were highly variable across sections, with no section displaying consistently higher or lower ranges (**Figure 2**). The highest average percentage of grid cells occupied by a section with more than one species was section *Circumdati* (n = 12 species), with an average of 2.72% grid cells occupied. Section *Aenei* had the lowest average (1.81% grid cells occupied; n = 7 species). Examination of species richness maps by section showed that many sections have similar distributions as the entire genus (**Figure S10**). However, the distributions of a few sections differ from the rest. For example, species in section *Sparsi* (n = 7) are restricted to most of Central and South America, southeastern Asia, and central Africa, whereas species in section *Restricti* (n = 13 species) are mainly predicted to be absent from Africa.

**Figure 2:**
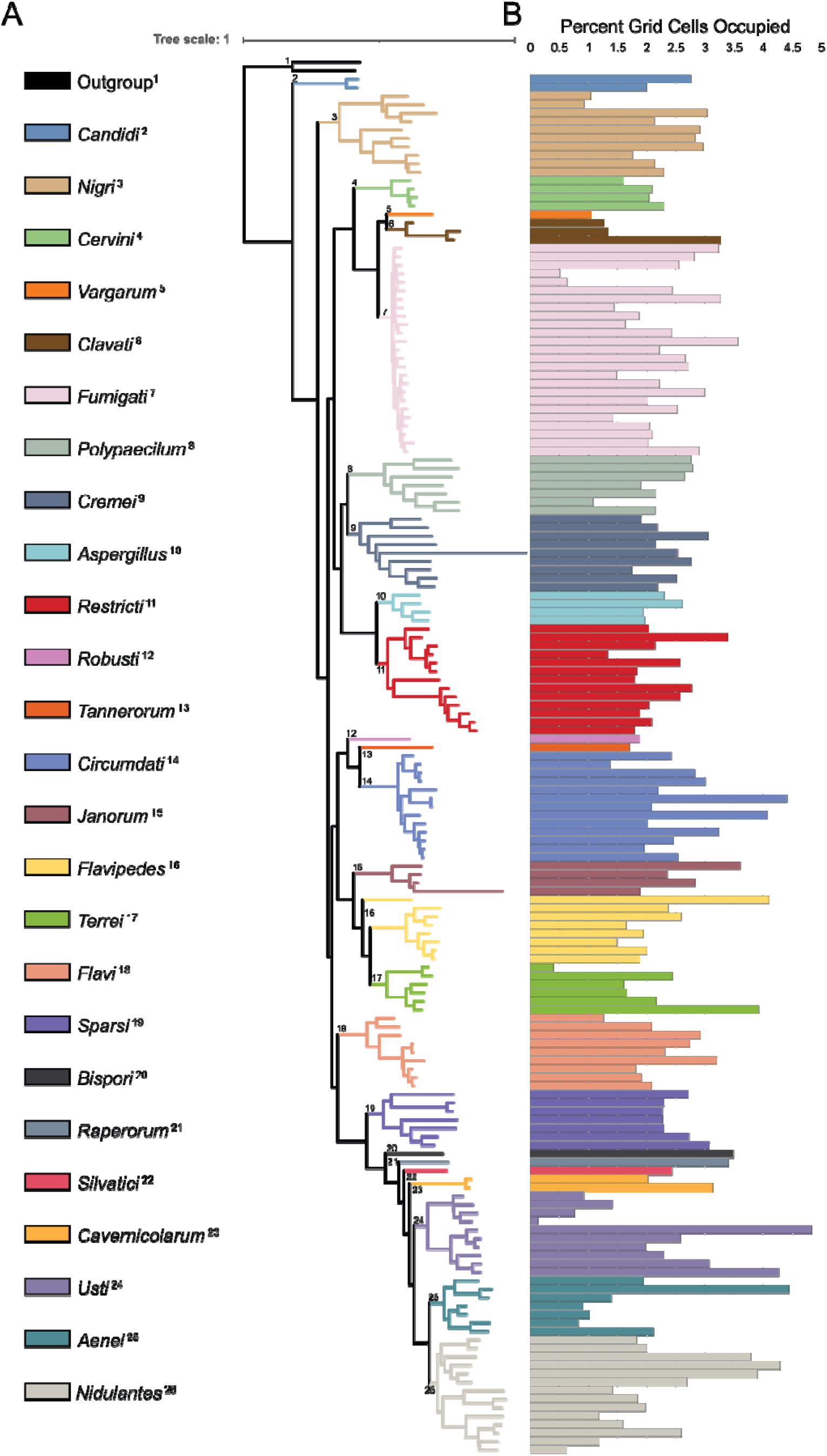
Species ranges vary extensively within taxonomic sections and across the genus. A) Phylogeny of 163 *Aspergillus* species. Colored branches and numbers on the phylogeny correspond to taxonomic sections, which are depicted on the left. The species *Talaromyces mimosinus* and *Talaromyces marneffei* were used as the outgroup and are designated by the black branches at the top of the phylogeny. Each tip on the phylogeny corresponds to a species. Species names have been removed for easier visualization but can be viewed in **Figure S8**. B) Species ranges represented by the percentage of grid cells (pixels) occupied. The X-axis shows the percent grid cells occupied by each species as a proxy for range, and the Y-axis is the phylogeny.

Previous studies in yeast have shown that species ranges are negatively correlated with species richness and absolute latitude (30). To test whether these patterns also held true in *Aspergillus*, we analyzed the relationship between range size and absolute latitude and species richness using phylogenetic generalized least squares analyses. Species range was statistically significantly negatively correlated with absolute latitude (p = 0.00002, effect size = −0.0244, r^2^ = 0.106, n = 163) (**Figure S9B** and **Table S5**). Species range was non-statistically significantly negatively correlated with average species richness per each species range (p = 0.169, effect size = −0.008, r^2^ = 0.012, n = 163) (**Figure S9C** and **Table S5**). Lastly, we did not find that phylogenetic distance and geographic distance were correlated (Mantel test based on Pearson Product-Moment, p = 0.35; Mantel test statistic = 0.0107; n = 163 species) (**Figure S11** and **Table S5**).

### Few species have large proportions of overlap in ranges

As many of the species in this analysis occupy similar regions, we sought to quantify the amount of geographic overlap between species by calculating the pairwise Jaccard Index of similarity (**Figure S12** and **Table S6**). Jaccard Index values closer to one indicate identical ranges, where values of zero indicate no overlap. The species pair with the largest amount of overlap was *Aspergillus puniceus* (section *Usti*) and *Aspergillus neoniveus* (section *Flavipedes*) with a Jaccard index of 0.79, indicating a substantial portion of their ranges overlap (**Figure 3A**). In contrast, *Aspergillus leporis* (section *Flavi*) did not overlap with *Aspergillus puniceus* (section *Usti*) or *Aspergillus pulvinus* (section *Cremei*) (**Figure 3B**). Across all 176 species, most of them display low amounts of overlap in ranges indicated by an average global species richness of ∼29 species / per grid cell, but the extent of the overlap is variable. Most species pairs had Jaccard Indices between 0.1 and 0.3, and very few were greater than 0.7 (**Figure 3C**). These results suggest that while many species do overlap in some areas, the overlap in ranges between species pairs is relatively small, resulting in low average global species richness. The low amount of overlap could be attributed to competitive exclusion, niche differentiation, or environmental partitioning, with similar species sharing similar ecological niches and competing for limited resources and environmental conditions, which could result in reduced co-occupation of environments. Additionally, there could also be differences in microhabitats, with variations in soil types, climate, or vegetation types, which enable the formation of distinct ecological niches that may allow for species to occupy similar regions without necessarily overlapping in ranges.

**Figure 3:**
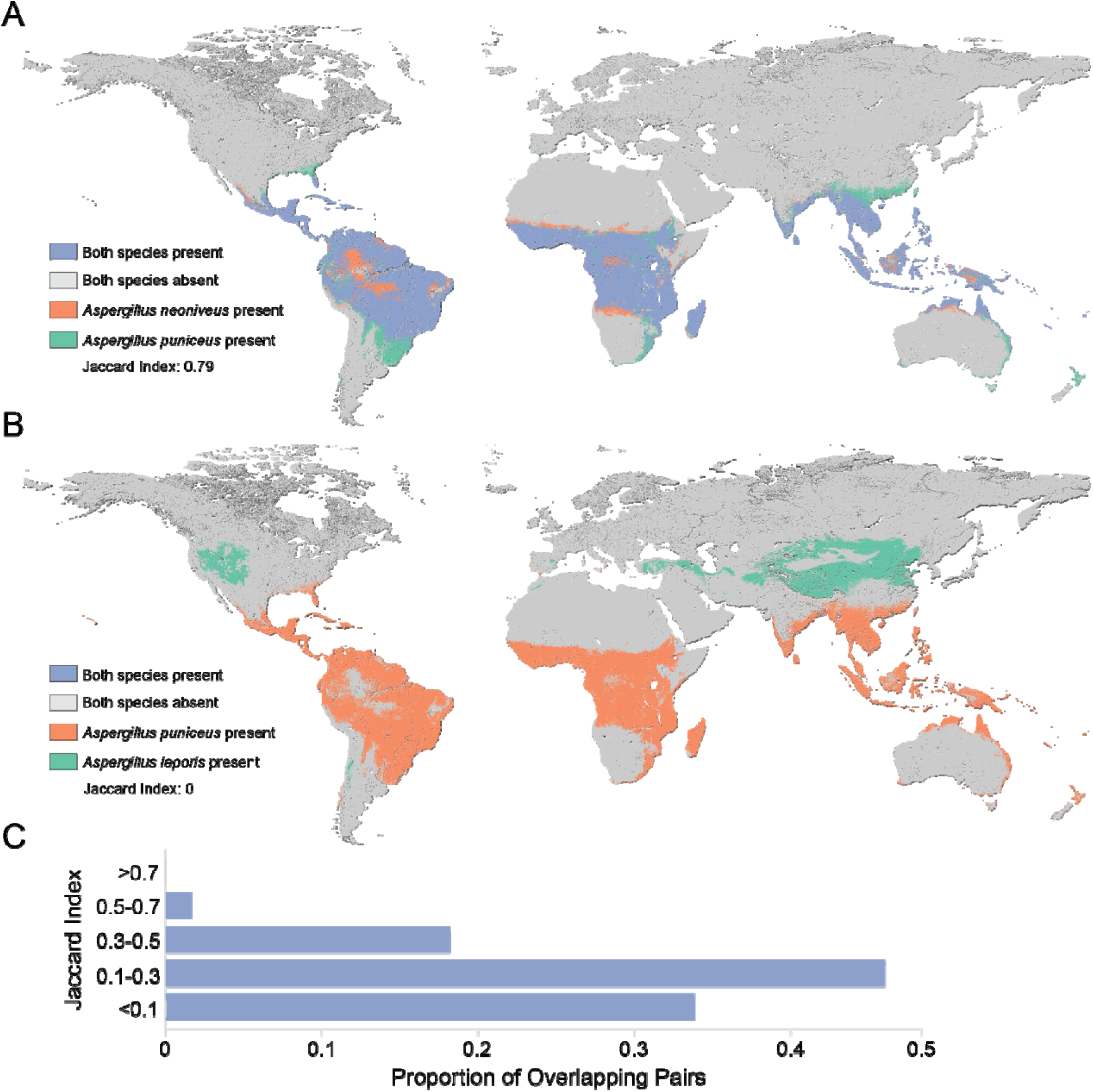
Most species pairs exhibit limited overlap in their ranges with a few exceptions. Purple represents areas where both species have predicted presence values, grey are areas where both species have predicted absence values, and orange and green are areas where only one species is present. Jaccard Index is displayed for both species pairs with values of 1 indicating identical ranges and 0 indicating no overlap. A) *Aspergillus neoniveus* (section *Flavipedes*) and *Aspergillus puniceus* (section *Usti*) have the largest proportion of range overlap even though each species individually does not have the overall largest range. The Jaccard Index of overlap for this species pair is 0.79. B) *Aspergillus puniceus* (section *Usti*) and *Aspergillus leporis* (section *Flavi*) do not overlap at all and have a Jaccard Index of 0. *Aspergillus leporis* generally occupies northern latitudes, whereas *Aspergillus puniceus* generally occupies southern latitudes. C) Most species pairs share limited overlap in range, with the rare occurrence of large range overlaps and an average species richness of ∼29. X-axis indicates the proportion of species pairs out of 15,400 unique combinations of pairs across 176 species. Y-axis shows the Jaccard Index bins. 0.34 species pairs overlapped with a Jaccard Index of <0.1, 0.46 between 0.1-0.3, 0.18 between 0.3-0.5, 0.02 between 0.5-0.7, and 0.0004 >0.7.

### Diverse environmental drivers of Aspergillus distributions

To better understand environmental factors that influence *Aspergillus* macroecology, we used SHapley Additive exPlanations (SHAP) values, which provide a quantitative measure of the contribution of each feature to the random forest classifier’s prediction for individual species. Across all 176 species, the most informative variables were Köppen-Geiger Climate classifications, ecofloristic zones, index of cumulative human impact, soil class, and biome (**Figure S2** and **Table S3**). Four of the top five predictors (Köppen-Geiger Climate classifications, ecofloristic zones, soil class, and biome) are categorical and reflect diverse environmental conditions. This suggests that variations in climate zones, vegetation types, and soil characteristics are key in determining species ranges. The index of cumulative human impact in the top predictors highlights the importance of anthropogenic factors. This index, which aggregates various human activities such as land use, population density, and overall human infrastructure, reveals that *Aspergillus* species may thrive or decline in response to human influence on their habitats. This suggests that *Aspergillus* species distributions are also driven by human activities that alter ecosystems or create more available niches. Considered in combination, these five variables suggest that *Aspergillus* species distributions are shaped by both natural environmental gradients and human-induced changes.

To further explore which variables were the most predictive of average species richness per ecoregion, we conducted negative binomial regressions, scaled linear regressions, and a relative importance analysis on 95 variables (88 quantitative variables and 7 binary variables). To reduce correlations between significant variables, we decomposed highly correlated variables into principal components, which yielded 47 variables or principal components (**Figure S13** and **Table S9**). We found Index of Cumulative Human Impact, Temperate vs. Non-Temperate ecoregions, Productivity PCA, and Temperature PCA were the most predictive variables (relative importance value of 1), followed by Clay PCA and Forest vs. Non-Forest ecoregions (both with a relative importance of 0.99) (**Figure 4** and **Table S11**). Consistent with the random forest results, we found that both environmental characteristics and human activities influence the distribution of *Aspergillus* species (**Figure 4**).

**Figure 4:**
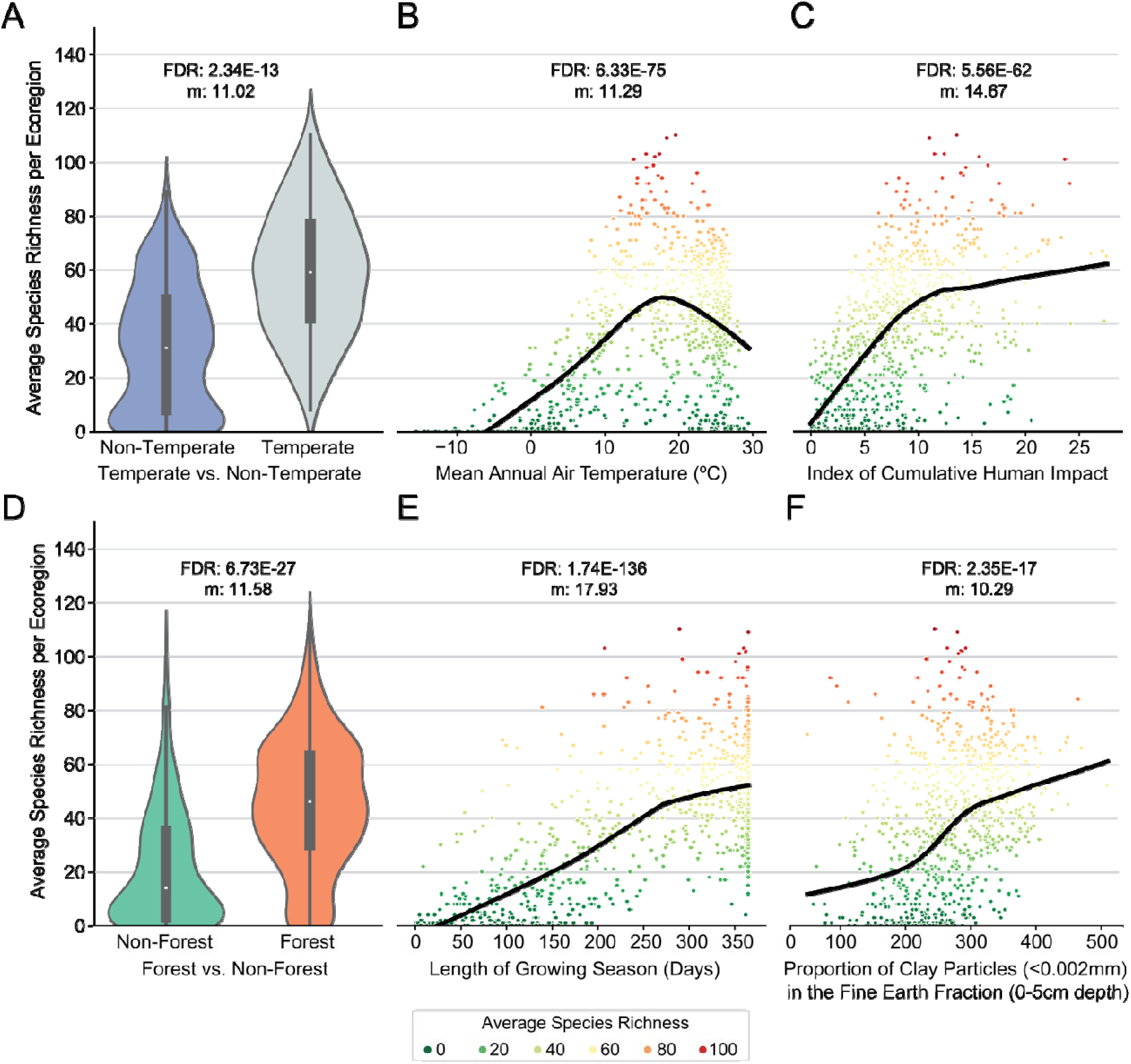
The six environmental variables most predictive of *Aspergillus* average species richness per ecoregion. The negative binomial regressions and scaled linear regressions of the 47 environmental variables and/or principal components revealed the six most predictive variables of average species richness per ecoregion. A-C) The Y-axis displays the average species richness per ecoregion across three environmental variables: (A) Temperate vs. Non-Temperate ecoregions, (B) Mean Annual Air Temperature (°C), which had the largest effect size from the Temperature PCA (FDR: 1.54E-88, m: 10.2), and (C) Index of Cumulative Human Impact. D-F) The Y-axis displays the average species richness per ecoregion across three environmental variables: (D) Forest vs. Non-Forest ecoregions, (E) Length of the Growing Season (Days), which had the largest effect size from the Productivity PCA (FDR: 1.70E-112, m: 16.34), and (F) Proportion of Clay Particles in the Fine Earth Fraction which had the largest effect size from the Clay PCA (FDR: 1.15E-17, m: 10.24). Each plot contains the False Discovery Rate (FDR) of the negative binomial regression, scaled slope of linear regression (m), and black lines represent the locally estimated scatter plot smoothing. Each dot on the scatter plots corresponds to an ecoregion, and the colors relate to average species richness, with warm colors indicating higher average species richness and colder colors indicating lower average species richness.

Latitudinal biodiversity gradients where species richness is typically higher in the tropics have been found across mammals, plants, invertebrates, and other microbes (33–37). Contrary to these analyses, we found that *Aspergillus* average species richness was higher in temperate ecoregions than in tropical ones (**Figure 4A** and **Figure S14**). This result aligns with findings from other fungal biodiversity studies (19,30,38). The Temperature PCA suggests that regions with average temperatures between ∼10°C and ∼25°C support the highest species richness (**Figure 4B**). Additionally, the Productivity and Clay PCAs emphasize the role of ecosystem productivity and clay content in soil in shaping *Aspergillus* distributions.

### Aspergillus species display variability in predicted responses to changing climates

To predict how *Aspergillus* species would respond to future climate change scenarios, we refined our random forest classifier, focusing on 15 environmental variables that have corresponding data under predicted climate change models (**Table S12**). We trained the classifier on the current dataset and used the trained classifier to predict species distributions across two future timeframes: 2041-2070 and 2071-2100. There were three predicted distributions per timeframe reflecting three distinct climate change scenarios: mild (sustainability, respect of environmental boundaries, and lower resource and energy intensity), moderate (regional rivalry redirecting focus to national and regional security, environmental concerns are low priority resulting in strong environmental degradation in some regions), and severe (fossil-fueled development, exploitation of fossil fuels to increase development and growth of the global economy) (39,40). We predicted distributions for species with > 300 occurrence records and excluded species with True Positive and True Negative rates < 75%, which yielded predictions for 33 species (**Figure S15** and **Table S13**). We compared the predictions of the current distributions of species using the original vs. the refined classifier and found that the refined classifier predicts larger ranges for species in the same locations (average of ∼ 0.5% larger), which is likely a result of focusing solely on climatic factors.

Our classifiers predicted that there was an overall range decrease for many *Aspergillus* species across all timeframes and climate change scenarios (**Figure S16**). However, we found substantial variation in the magnitude and direction of changes, highlighting the nuanced interactions between climate change and individual species ranges (**Figure 5 and Figure S16**). Specifically, 14 species, including *Aspergillus versicolor* (section *Nidulantes*), *Aspergillus ochraceus* (section *Ochraceorosei*), and *Aspergillus terreus* (section *Terrei*) were predicted to expand their ranges due to climate change, although some expansions appear marginal or suggest more stable distributions, as seen with *Aspergillus niger* (section *Nigri*). Species from sections *Nigri* and *Nidulantes* showed a higher tendency toward range expansions, raising the hypothesis that they may harbor specific evolutionary or ecological traits that make them more resilient or adaptable to changing climatic conditions. In contrast, *Aspergillus udagawae* (section *Fumigati*), a minor pathogen (41), was predicted to significantly contract its range, particularly in southern Africa and Central America.

**Figure 5:**
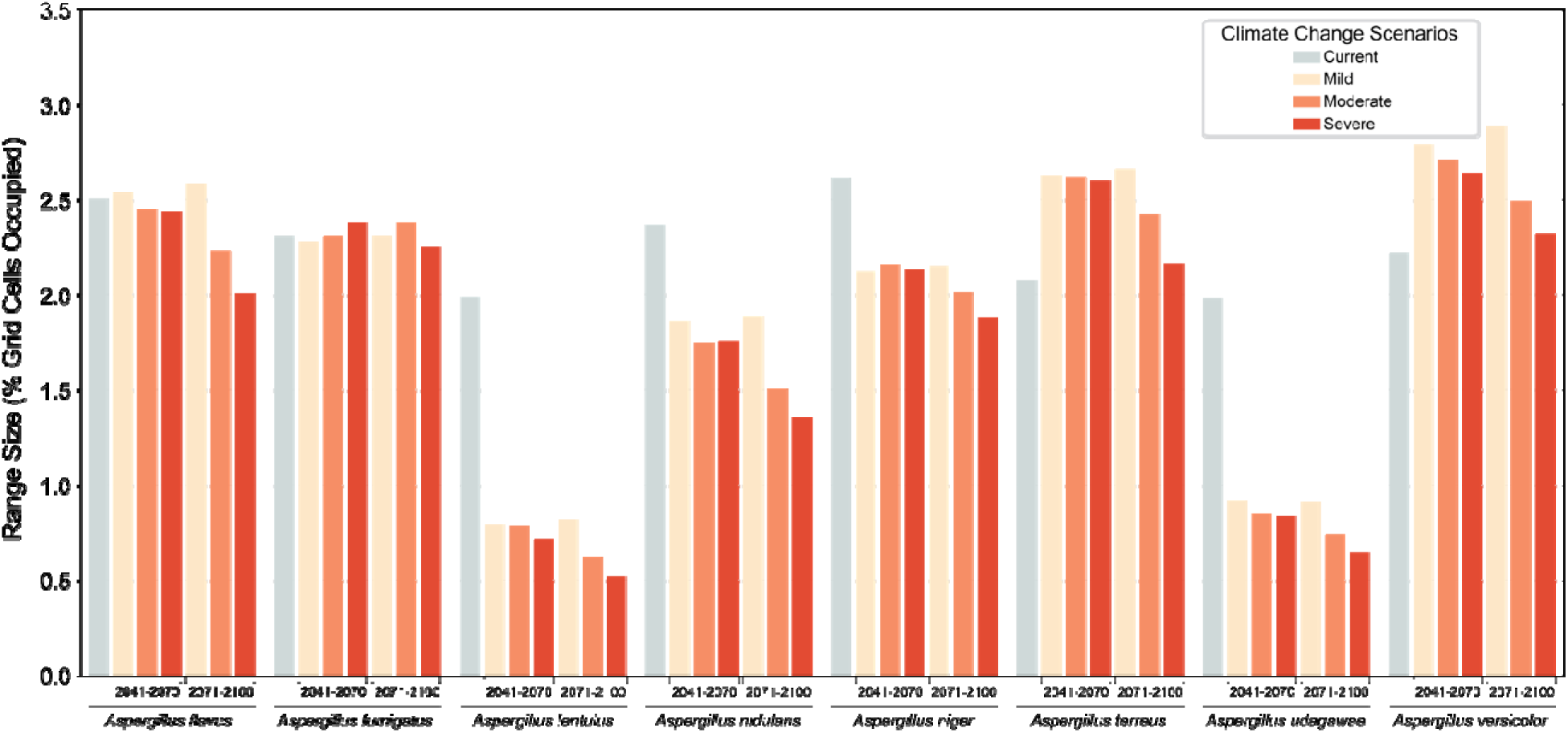
Some *Aspergillus* species ranges are predicted to increase and some to decrease under warming temperatures. Different species have varying changes in their predicted ranges under future climate change scenarios. X-axis shows each species under the mild (sustainability, respect of environmental boundaries, and lower resource and energy intensity), moderate (regional rivalry redirecting focus to national and regional security, environmental concerns are low priority resulting in strong environmental degradation in some regions), and severe (fossil-fueled development, exploitation of fossil fuels to increase development and growth of the global economy) climate change scenarios for the two timeframes of 2041-2071 and 2071-2100. The Y-axis shows predicted ranges in percent grid cells (pixels) occupied. The grey bar is the predicted range using current data, the purple using mild scenario data, the orange bar using the moderate scenario data, and the green using severe scenario data. Some species have predicted range increases across timeframes and models, like *A. versicolor*. However, some have more variable responses (*A. terreus*) or overall decreases in range (*A. udagawae*).

A few species have overall predicted range increases from the second timeframe to the third (*Aspergillus caninus*, *Aspergillus ochraceus*, *Aspergillus puulaauensis*, and *Aspergillus ruber*). These predicted range increases correspond to a reduction in areas like southern Africa with much warmer forecasted air temperatures from current to 2041-2070 under the severe scenario and an increase in prevalence in eastern Asia.

The variation in predicted species ranges suggests that climate change will not necessarily lead to a global increase in the range of *Aspergillus* species. In some regions, urbanization or localized climate change could result in higher-than-average temperature increases, driving species expansions in urban heat islands or areas of intense land use change. Some species may exploit anthropogenically-altered habitats due to agriculture or infrastructure development. Additionally, many *Aspergillus* species are temperature tolerant and can grow in temperatures ranging from ∼ 10°C to ∼ 45°C, and in some species even higher (42,43). In this context, the lack of major predicted range expansions for most species has biological relevance, as a few degrees of temperature increase will likely not result in major changes in ranges. Furthermore, our observation that the refined classifier tends to predict larger ranges for most species, coupled with our finding that some species ranges remain stable, could suggest that climatic factors play a more significant role in predicting individual species ranges. In contrast, species that had the larger deviations between the classifiers may be more strongly influenced by environmental factors.

### The ranges of opportunistic pathogens tend to decrease over time

Rising temperatures due to climate change have been hypothesized to increase the ranges of fungal species, as environments with temperatures previously outside of optimal growth ranges become more habitable with increasing temperatures (44,45). Numerous species have observed range increases potentially in response to changing climates, including species in the genera *Blastomyces*, *Histoplasma*, and *Coccidioides*, which all have generally geographically restricted ranges (45–47). To determine the potential distributions of *Aspergillus* pathogens under the severe climate change scenario, we focused on two major pathogens (*Aspergillus fumigatus* and *Aspergillus flavus*) and six minor pathogens (*Aspergillus felis*, *Aspergillus lentulus*, *Aspergillus nidulans*, *Aspergillus niger*, *Aspergillus terreus*, and *Aspergillus udagawae*) from four different sections.

We found that the ranges of major pathogens are predicted to decrease slightly over time under the severe climate scenario (**Figure 6A, 6B, 6C**), particularly by the 2071-2100 period. While their combined ranges are predicted to decrease, *Aspergillus flavus* is predicted to initially increase in 2041-2070 and decrease in 2071-2100 to a similar level as the current range, and *Aspergillus fumigatus* decreases slightly but remains rather constant. However, the predicted changes are not uniform across all regions. During the 2041-2070 timeframe, range contraction is predicted in southern Africa and expansion in South America. This suggests that climate-driven range shifts for the major pathogens may be region-specific, driven by localized environmental variables, rather than on a global scale. The expansion of these pathogens in South America indicates that while species may experience geographic shifts, the overall terrestrial space they occupy may remain relatively similar (**Figure 6D**). In contrast, minor pathogens show a consistent decline in their predicted ranges over time (**Figure 6E, 6F, 6G, 6H**). There is a particularly large, predicted range contraction of minor pathogens in the United States and southern Africa, accompanied by a smaller but notable expansion in eastern regions of South America. While the ranges of these minor pathogens are predicted to decrease over time, they persist globally in relatively the same regions with variable prevalence.

**Figure 6:**
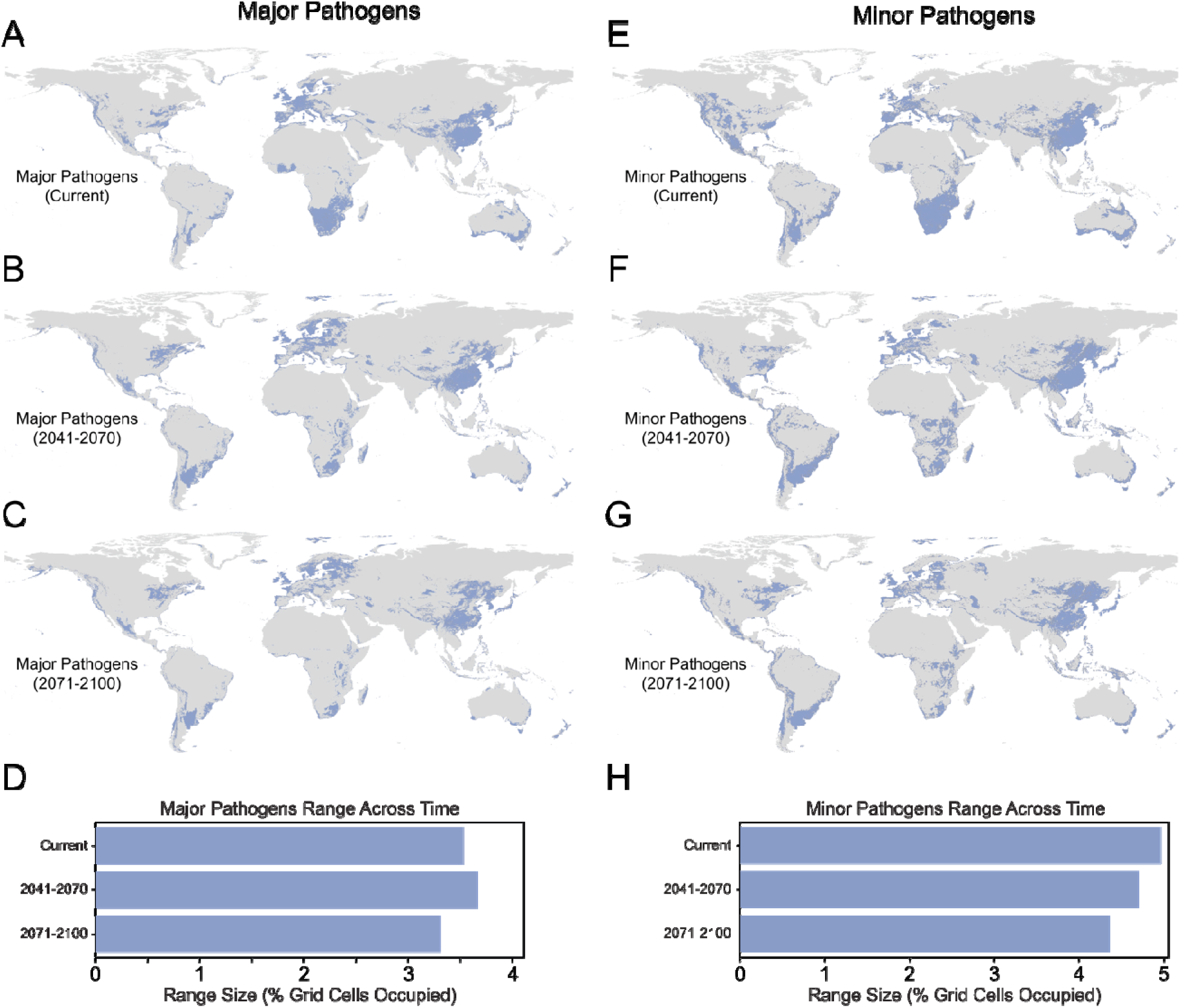
Prevalence of major and minor pathogens decreases under the severe or “worst-case scenario” climate change model. The maps depict the predicted distributions of major pathogens (*A. fumigatus* and *A. flavus*; left side) and minor pathogens (*A. felis*, *A. lentulus*, *A. nidulans*, *A. niger*, *A. terreus*, and *A. udagawae*; right side) under the severe climate model. This model is based on fossil-fueled development and an energy-intensive lifestyle. A-D) Major pathogen ranges for the timeframes: current (A), 2041-2071 (B), and 2071-2100 (C) and their ranges over time (D). X-axis is the range size in percent grid cells occupied (pixels) and the Y-axis represents the timeframes. E-H) Minor pathogen ranges for the timeframes: current (E), 2041-2071 (F), and 2071-2100 (G), and their ranges over time (H). The X-axis is the range size in percent grid cells occupied, and the Y-axis represents the timeframes. The presence of species is depicted in purple, and the absence is depicted in grey.

The observed variation in *Aspergillus* species ranges predicted under future climate change scenarios suggests that species are expanding in regions where they already reside or being eliminated in regions where they can no longer inhabit due to changing climate. We analyzed species richness across timeframes and climate change scenarios and found that most areas have an overall reduction in species richness apart from eastern Asia and South America, where species richness appears to increase under the moderate and severe scenarios (**Figure S17**). *Aspergillus* species have been noted as generalists, and it is thought that generalist species may be more resilient to changing climates because of their ability to occupy a wider diversity of environments (3). This generalist lifestyle, coupled with their global presence, may explain their ability to withstand changing climate conditions. We have shown *Aspergillus* species display complex and diverse ecologies that are not entirely reliant on climatic conditions, which indicates that warming environments will not necessarily lead to a higher abundance of species.

## Conclusions

In this study, we investigated patterns in the global distributions of *Aspergillus* species. We used a combination of species range modeling and environmental variables to generate a comprehensive picture of species in the genus *Aspergillus* across diverse environments. Our results revealed that species richness is highest in temperate forested regions with stable, warm, humid climates, particularly Mediterranean-type ecosystems. We also identified five climatic and environmental factors that are the most predictive of average species richness and found that human activities may have a crucial role in shaping *Aspergillus* distributions. Interestingly, we observed deviations from traditional patterns in plants and animals while analyzing *Aspergillus* distributions, such as a lack of latitudinal and longitudinal gradients and higher species richness in temperate regions compared to tropical regions. Additionally, we identified extensive variability in the species range sizes within and across sections, and a small proportion of species ranges overlap, showing that some species are more widespread while others are predicted to have restricted, localized ranges. Lastly, we found *Aspergillus* species have highly variable responses to future climate change scenarios, depicting the species-specific changes as some increase, decrease, or remain rather stable.

One key component to consider while interpreting these results is that these predictions assume that species will continue to occupy the same ecological niches. If species evolve (e.g., through mutations that facilitate growth at higher temperatures) in response to shifting environmental conditions, particularly in regions with rapid temperature increases or other climate-related stresses, their future distributions may diverge extensively from these predictions. Experimental evolution studies in various fungi have shown that the acquisition of mutations that enable thermotolerance can occur in relatively short time frames (48–52); thus, increasing global temperatures and potential range shifts or expansions of fungal species have led to concerns that adaptation to these increasing temperatures will result in the emergence of new opportunistic pathogens of humans (23–25,53). The evolution of thermotolerance may allow fungi to more readily infect humans (23–25), but whether a few degrees in temperature increase will give rise to new fungal pathogens remains uncertain (43). These predictions may aid in determining which species may maintain their presence in areas where mean annual air temperatures meet or exceed human body temperatures to help monitor potential emerging *Aspergillus* pathogens.

## Methods

### Retrieval and pre-processing of species occurrence records

All occurrence records (116,664) were obtained from the GlobalFungi database (22) (release 5, January 8^th^, 2024) and preprocessed following the protocols of a previous study (30). Briefly, we selected only valid species names listed Index Fungorum and reconciled species names with the updated nomenclature. We removed duplicate coordinates for the same species. Then we used the R package CoordinateCleaner (54) to remove records with equal longitude and latitude coordinates, zero coordinates, and coordinates that matched the centroid of counties/provinces or biodiversity institutions. This resulted in 41,820 occurrence records from 240 *Aspergillus* species.

Next, for each occurrence record, we extracted data from 96 environmental raster files (30) that were then projected onto the WGS84 (EPSG:4326) coordinate system with a resolution of 30” (∼1km^2^) using a custom R script with the terra and raster libraries (55,56). The raster files contained information about the vegetation, soil, climate, and anthropogenic inputs of the area (**Table S1**) (30). Following the environmental variable extraction, we removed occurrence records with NA values and records with the same environmental variables within a hundredth-degree latitude or longitude to prevent the overrepresentation of samples from a specific location. This resulted in a training dataset of 30,542 occurrence records across 236 species. In addition to the occurrence records, we randomly sampled coordinates across the entire raster extent (102,609 points) as pseudoabsence points.

### Training the random forest algorithm

Using the data compiled for *Aspergillus* species, we trained a random forest classifier to predict species distributions. We used the same parameters and methods as the study of David et al. (30). Briefly, we used the R package randomforest (57) with a downsampling approach to reduce overfitting. We created a model for each species with at least five occurrence records, which resulted in 205 random forest classifiers (one for each species) (**Table S2**). Each classifier had 100 decision trees, and all other parameters were set to their default values. We used a leave-one-out strategy for validation, which consists of leaving out one row from each training dataset for validation. We retained 176 models with at least a 75% True Positive rate and Negative rate for further analysis. The average area under the curve for the Receiver Operator Characteristic curves for all 205 models was 0.93, and an average True Positive rate of 85% and True Negative rate of 90%. The summary statistics from the random forest predictions for all 205 models can be found in **Tables S2 & 3** and **Figure S2**. We used the SHapley Additive exPlanations (SHAP) python package (58) to visualize and interpret feature importance from the random forest classifiers.

### Species range and overlap analyses

To estimate the ranges for each species, we calculated the number of grid cells (pixels) occupied per species from their predicted raster files using a custom R script. To determine if there were any meaningful differences in species ranges between the different taxonomic sections within *Aspergillus*, we built a concatenation-based maximum likelihood phylogeny using the nucleotide sequences Beta-Tubulin, Calmodulin, and RNA Polymerase Beta (second largest subunit) for 456 *Aspergillus* species obtained from previous studies (1,2). We included *Talaromyces marneffei* and *Talaromyces mimosinus* as outgroups, bringing the total to 458 species. Briefly, we created multi-fasta files for each gene, aligned the gene sequences across all 458 taxa using Mafft (v7.245) (59), built a concatenation matrix of the trimmed sequences using PhyKIT (v1.12.5) (60), and built the phylogeny using IQTree (v2.2.6) (61) (**Figure S1 and Table S14**). Only 163 of the 176 species in these analyses had nucleotide sequences available to analyze on the phylogeny. To display only the 163 species of interest, we pruned the phylogeny using the R Shiny application Treehouse (62).

To quantify the degree of overlap in range estimates between species, we calculated the Jaccard Index or similarity coefficient (total number of overlapping cells/total number of unique cells occupied by each species). Figures displaying species overlap in ranges were generated using QGIS.

To determine if there was a relationship between phylogenetic distance and geographic distance among species, we performed a Mantel test based on Pearson’s product-moment correlation. We identified the centroid of each species range using the terra package in R (55) and pruned the phylogeny to match the species labels using the R package ape (63). We calculated the pairwise geographic distances between species centroids using the Haversine formula using the geosphere package in R (64) and calculated the pairwise phylogenetic distances using the cophenetic function (the sum of the branch lengths that connect two species in the phylogeny) from the ape package. This analysis resulted in two distance matrices, which were aligned to each other by species. We used the resulting distance matrices to do the Mantel test using the vegan package in R (65) and evaluated the significance of the correlation using 999 permutations.

Additionally, we performed two phylogenetic generalized least squares (PGLS) analyses to test whether the range sizes of each species are influenced by absolute latitude and species richness. We extracted the centroid of each species range, determined the absolute latitude at the centroid of the species range, and used the R packages caper (66) and ape (63) to do the PGLS. In the same fashion, we extracted the average species richness across each individual species range and performed the PGLS.

### Assessing the ecological parameters that drive Aspergillus species diversity

To determine the ecological drivers of *Aspergillu*s diversity, we constructed negative binomial regression models (30) using 95 environmental variables (as the independent variable) and *Aspergillus* average species richness per ecoregion (as the dependent variable). We chose to look at the average species richness per ecoregion as compared to overall species richness or per biome because ecoregions represent more localized ecosystems with unique variations in climatic, geological, and biotic factors, and it is less computationally intensive. To do this, we selected 88 quantitative environmental variables from the random forest training data and extracted their average values for each ecoregion. In addition to the quantitative variables, we included 7 categorical variables (tropical, temperate, secondary forest, cultivated, continental, dry) encoded into binary variables based on the majority class of that variable in each ecoregion (temperate (1) vs. non-temperate (0) ecoregions). A species was considered present in that ecoregion if it was found in at least 10% of the region’s grid cells (pixels). We also performed scaled linear regressions with the slope (m) as a measure of effect size. We removed seven variables with False Discovery Rates (FDR) > 0.05 from the negative binomial regression. We then combined highly correlated variables into principal components to reduce correlations between environmental variables, which resulted in a total of 88 variables decomposed into 47 variables and/or principal components. Following decomposition, the greatest r^2^ between principal components/variables was 0.77 (µ=0.09) (**Figure S13**). The first PC for each principal component analysis explained at least 86% of the total variation apart from the soilRichness PCA (78%) (**Table S10**). There was an overall mean variance of 92%.

To determine the most predictive variables of average species richness out of all 47 variables and/or principal components, we used a relative importance analysis. We constructed negative binomial regression models using average species richness as the dependent variable for every combination of variables/ principal components whose linear relationship with average species richness had an r^2^ > 0.15 and slope (m) > 0.20. There were six variables and/or principal components that met the criteria: 2009 human footprint (index of cumulative human impact), temperate ecoregions, temperature PCA, productivity PCA, clay PCA, and forest ecoregions. This resulted in 63 individual models. We then calculated Akaike weights to estimate the relative importance of each variable.

### Predicting species distributions under climate change scenarios

To predict the future distributions of species under different climate change scenarios or shared socioeconomic pathways (SSPs), we selected 15 environmental variables from the larger dataset of 96 environmental variables that had future forecasts for the years 2041-2070 and 2071-2100. The future forecast data were obtained from the Climatologies at high resolution for the earth’s land surface area (CHELSA) CMIP6 database (67). To reduce the number of assumptions, only 15 environmental variables with future data were selected (**Table S12**).

For each of the 15 variables, raster files were obtained for both timeframes as well as three different SSPs: ssp126 (SSP1-RCP2.6 climate as simulated by the GCMs), ssp370 (SSP3-RCP7 climate as simulated by the GCMs), and ssp585 (SSP5-RCP8.5 climate as simulated by the GCMs) from the gfdl-esm4 models from the National Oceanic and Atmospheric Administration, Geophysical Fluid Dynamics Laboratory at a resolution of 30” (∼1km^2^) and the WGS84 (ESPG:4326) coordinate system. We classified the three SSPs as mild (ssp126, based on sustainability, respect of environmental boundaries, and lower resource and energy intensity), moderate (ssp370, regional rivalry redirecting focus to national and regional security, environmental concerns are low priority resulting in strong environmental degradation in some regions), and severe (ssp585, fossil-fueled development, exploitation of fossil fuels to increase development and growth of the global economy) climate change scenarios. We used the same occurrence records and filtering steps mentioned above. We created seven raster stacks: 1) current rasters (1981-2010) used in the predicting species distributions based on current data, 2) three raster stacks for the timeframe 2041-2070 for each climate change scenario, and 3) three raster stacks for the timeframe 2071-2100 for each climate change scenario. We then extracted data for each occurrence record and the pseudo-absence coordinates from the current climate raster stack. We filtered the training data in the same fashion as the previous model; however, with fewer variables, there were fewer NA values, resulting in 34,718 species occurrence records for training the model.

We trained the model on the current climate data to get their predicted distributions under current climatic and ecological conditions using the 15 environmental variables. However, in this case, we selected the 33 species with over 300 occurrence records to predict future distributions, which yielded 33 models. This resulted in an average area under the Receiver Operator Characteristic curve of 0.94. We then removed species with True Negative rates and True Positive rates less than 75% (**Figure S15**). All species met the criteria with an average True Positive rate of 84% and a True Negative rate of 89%. Once we had the current predicted distributions, we then used the trained model for each species to predict future distributions using the three different raster stacks for each timeframe with the assumption that species will occupy similar niches under the three forecasted climate change scenarios. This resulted in a total of seven predicted raster files per species. We then calculated the species ranges for each raster using the same approach as above and compared them to each other.

## Data Availability

All supplementary figures, data files, and code required to replicate the species distribution modeling and additional analyses have been deposited to the figshare repository and will be made publicly accessible upon publication.

## Financial Disclosure

This work was supported by the National Science Foundation (NSF Graduate Research Fellowship to OLR, DBI-2305612 to KTD, and DEB-2110404 to AR), the National Institutes of Health/National Institute of Allergy and Infectious Diseases (R01AI153356 to AR), and the Burroughs Wellcome Fund (to AR). The funders had no role in study design, data collection and analysis, decision to publish, or preparation of the manuscript.

## Competing interests

AR is a scientific consultant for LifeMine Therapeutics, Inc. All other authors have declared that no competing interests exist.

