## Supplementary Figures for "Global patterns of species diversity and distribution in the biomedically and biotechnologically important fungal genus *Aspergillus*"

**Supplemental Figures**

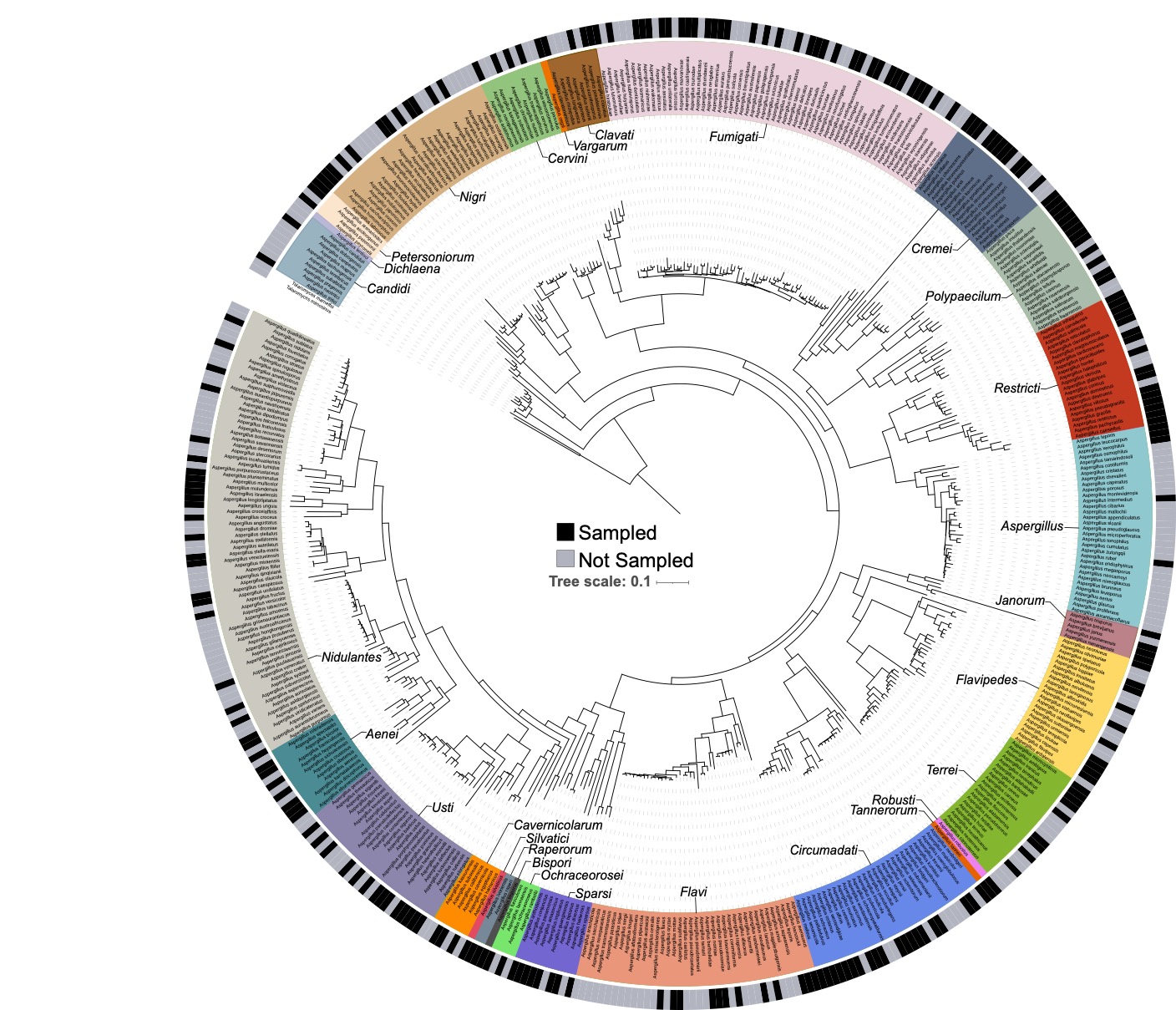

**Supplemental Figure 1:** Species geographic sampling across the genus *Aspergillus*. The large color blocks and labels indicate the 28 taxonomic sections. Black boxes indicate species that have been sampled from the environment. Grey boxes indicate species that lack sampling. 51 species were included in the analyses but were lacking sequences available to be included in the phylogeny.

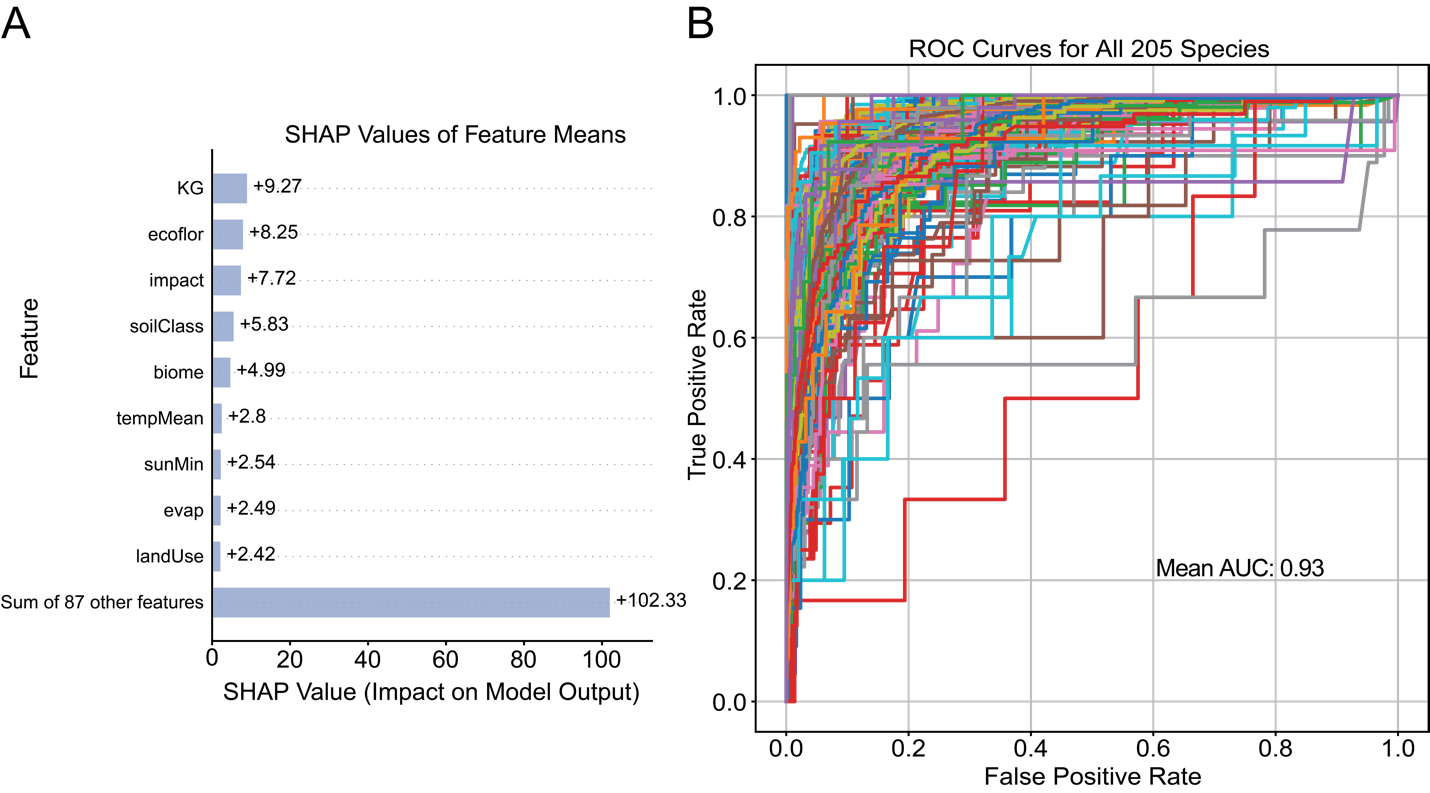

**Supplemental Figure 2:** A) Mean SHAP values for 96 features in random forest models and B) Receiver Operator Characteristic curves for all 205 species with greater than 4 occurrence records.

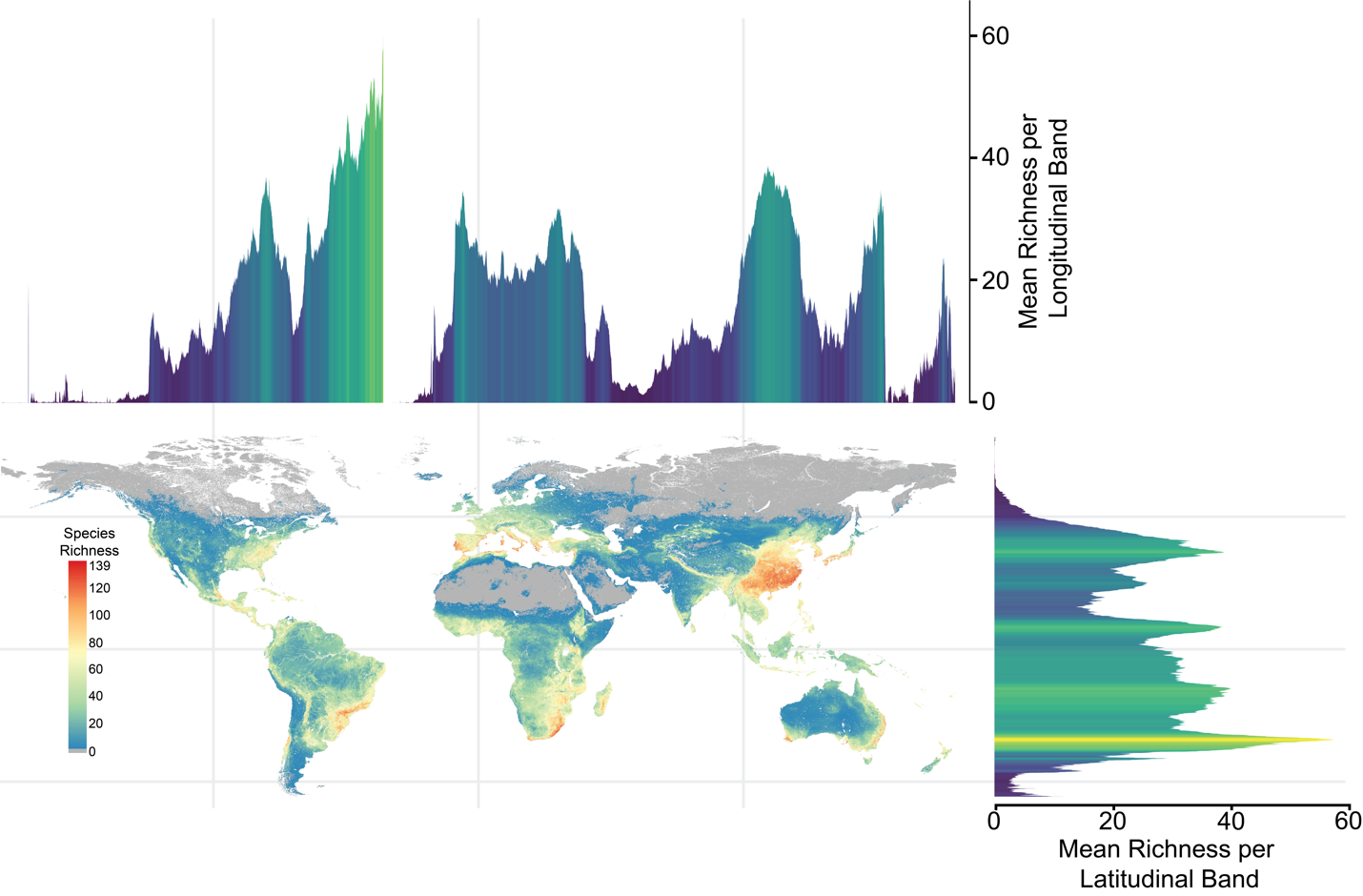

**Supplemental Figure 3:** Species richness and mean species richness per longitudinal and latitudinal bands.

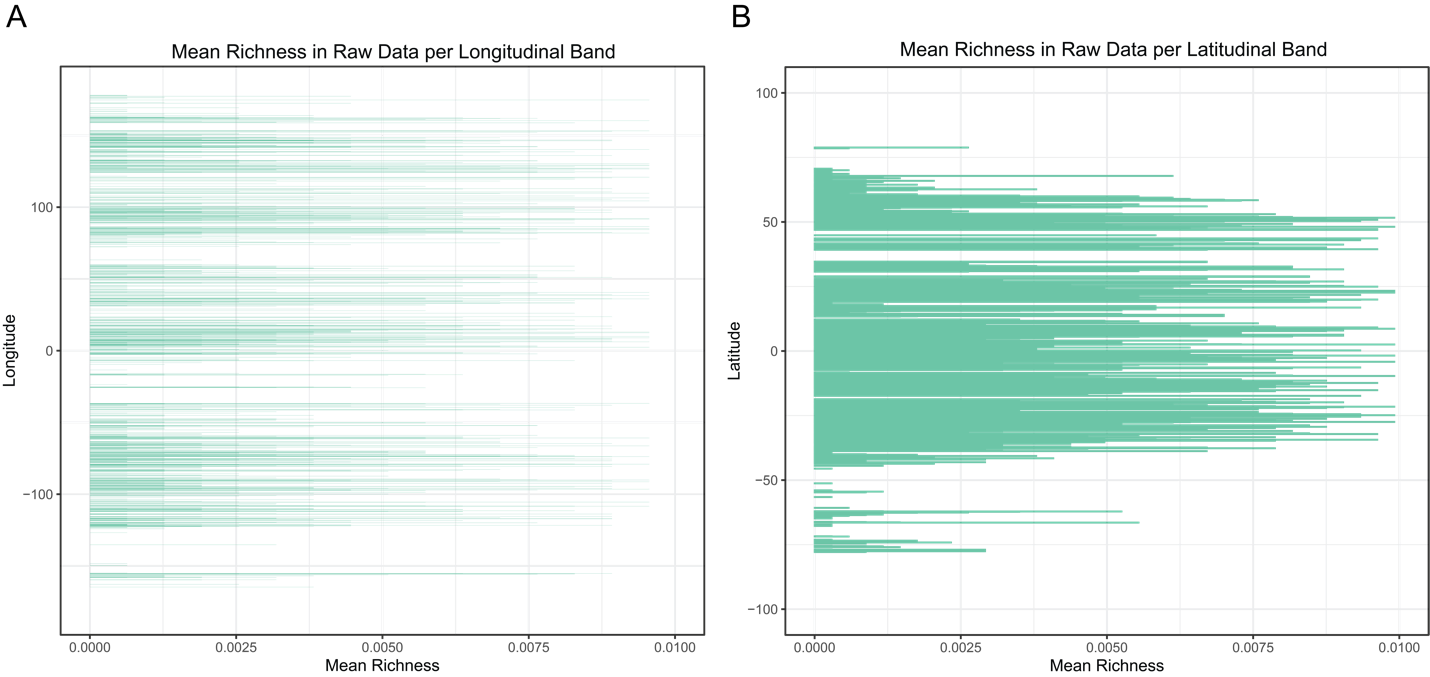

**Supplemental Figure 4:** No visible patterns in longitudinal and latitudinal richness in raw data from GlobalFungi. A) Mean richness per longitudinal band. B) Mean richness per latitudinal band.

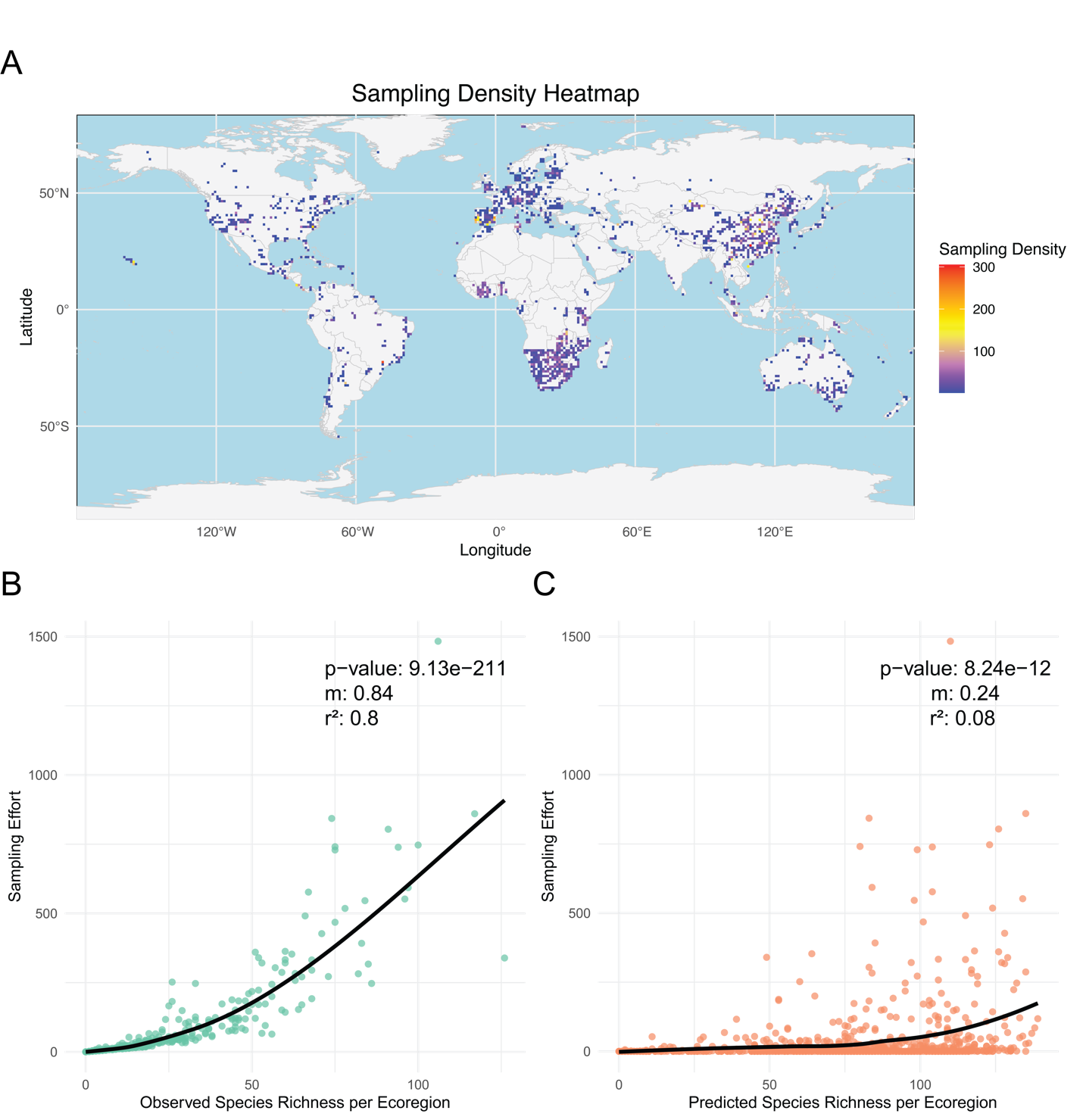

**Supplemental Figure 5:** Raw sampling data from GlobalFungi

A) Heat map of sampling density from the training data points for all species obtained from GlobalFungi. B) Sampling effort from the observation data in the training data from GlobalFungi for each ecoregion plotted against the observed species richness per ecoregion. C) Sampling effort from the raw data per ecoregion plotted against the predicted species richness per ecoregion. The p-value, scaled slope (m), and correlation coefficient (r^2^) of the linear model are also shown.

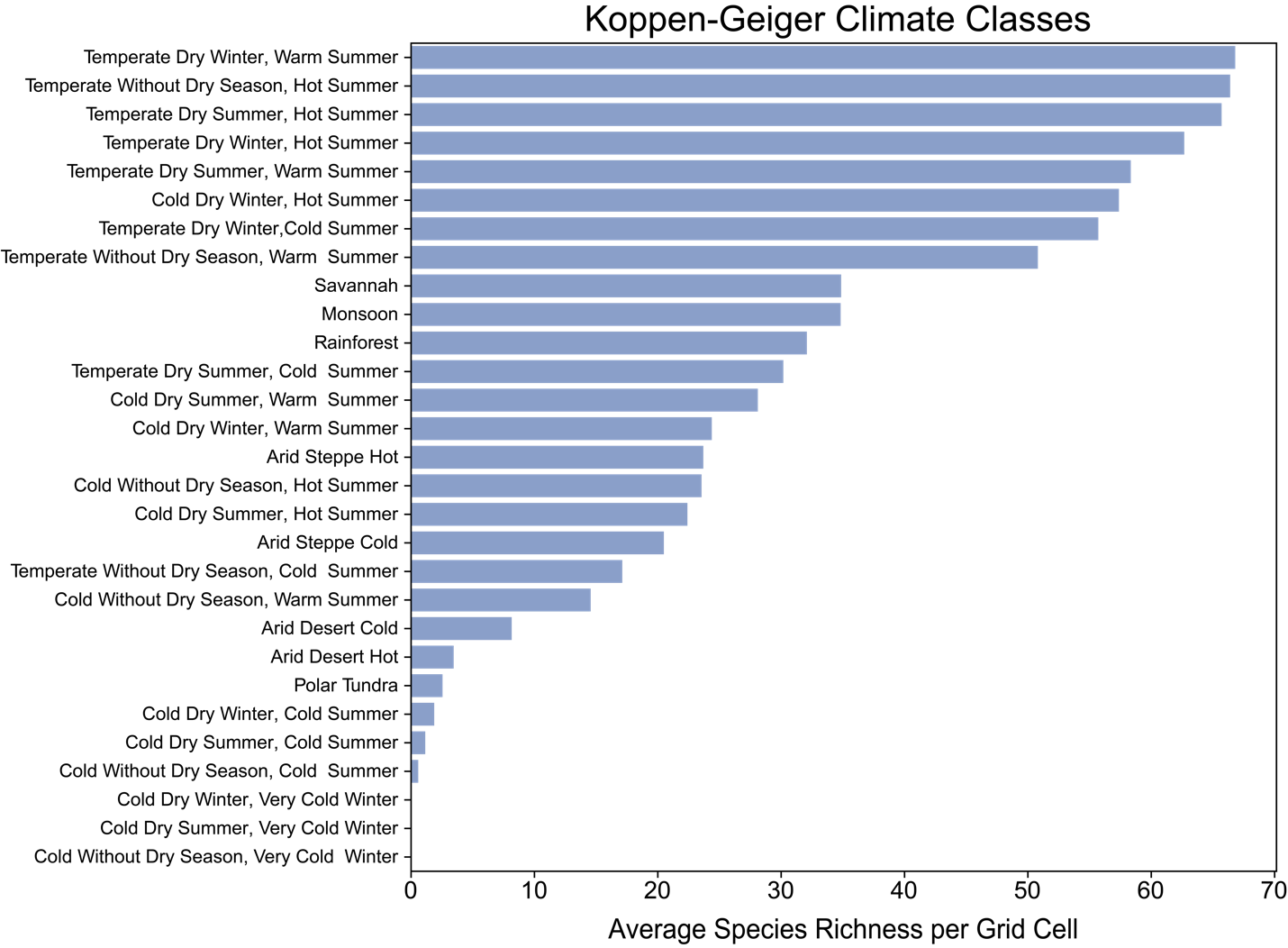

**Supplemental Figure 6:** Average species richness per grid cell (pixel) across Köppen-Geiger Climate classifications

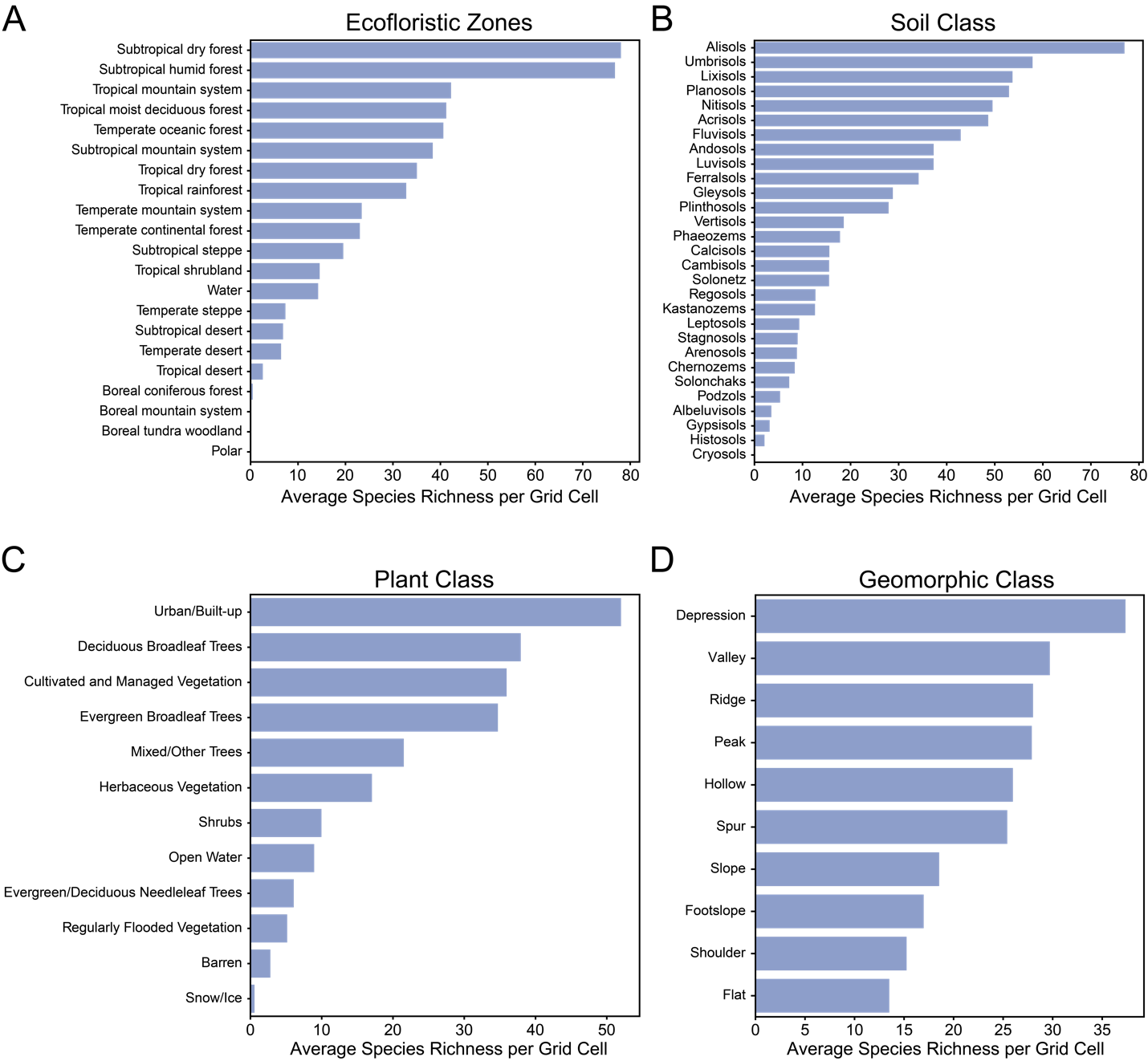

**Supplemental Figure 7:** Average species richness per grid cell (pixel) across A) Ecofloristic Zones, B) Soil Classes, C) Plant Classes, and D) Geomorphic Classes.

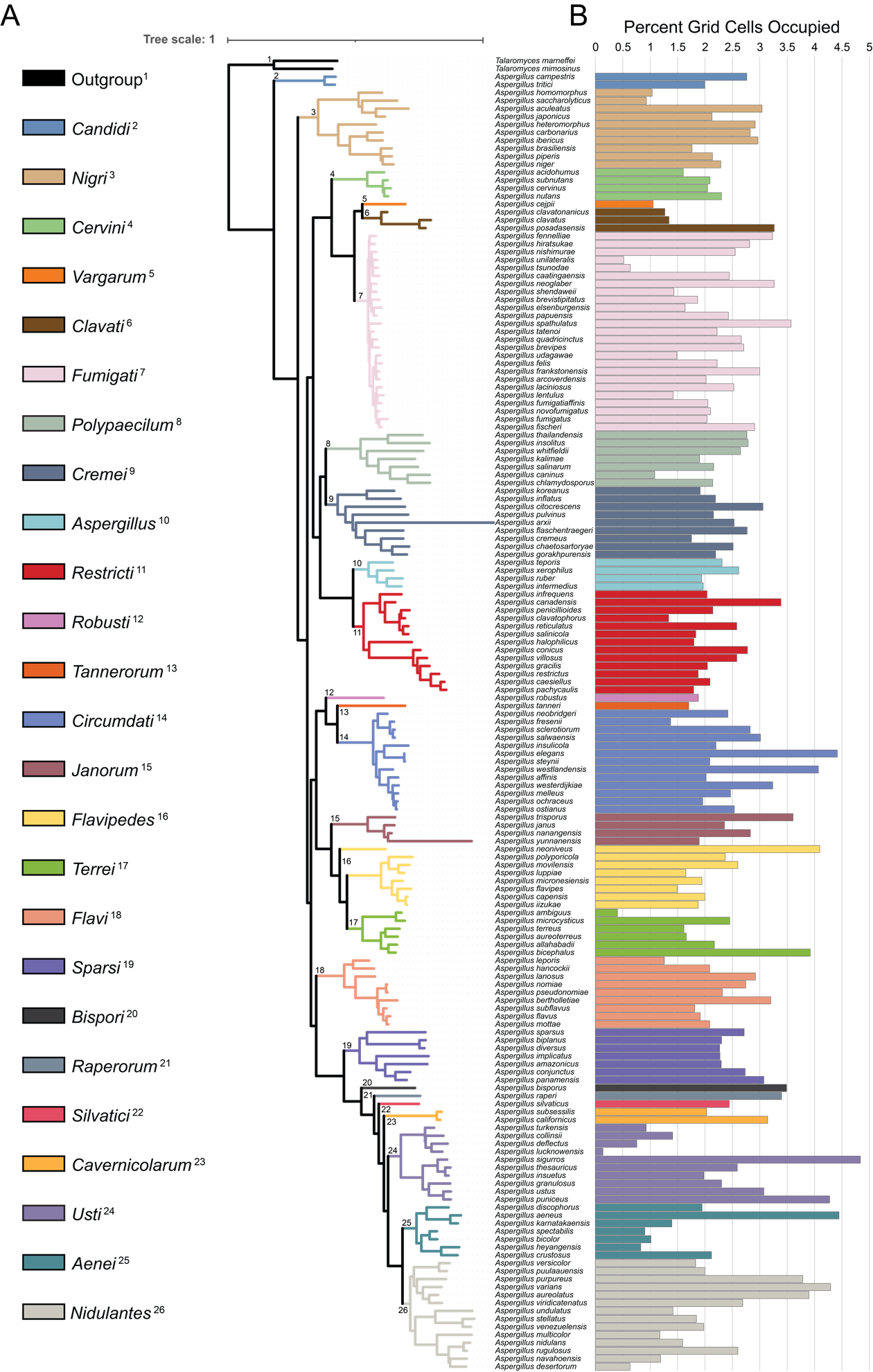

**Supplemental Figure 8:** Extensive variation in ranges across the genus and within taxonomic sections

A) Phylogeny of 163 *Aspergillus* species. Colored branches and numbers on the phylogeny indicate taxonomic sections corresponding to the colored boxes and numbers. The species *Talaromyces mimosine* and *Talaromyces marneffei* were used as the outgroup and are designated by the black branches. B) Species Ranges in percent grid cells (pixels) occupied. The X-axis displays species names and the Y-axis shows the percent grid cells (pixels) occupied by each species as a proxy for range.

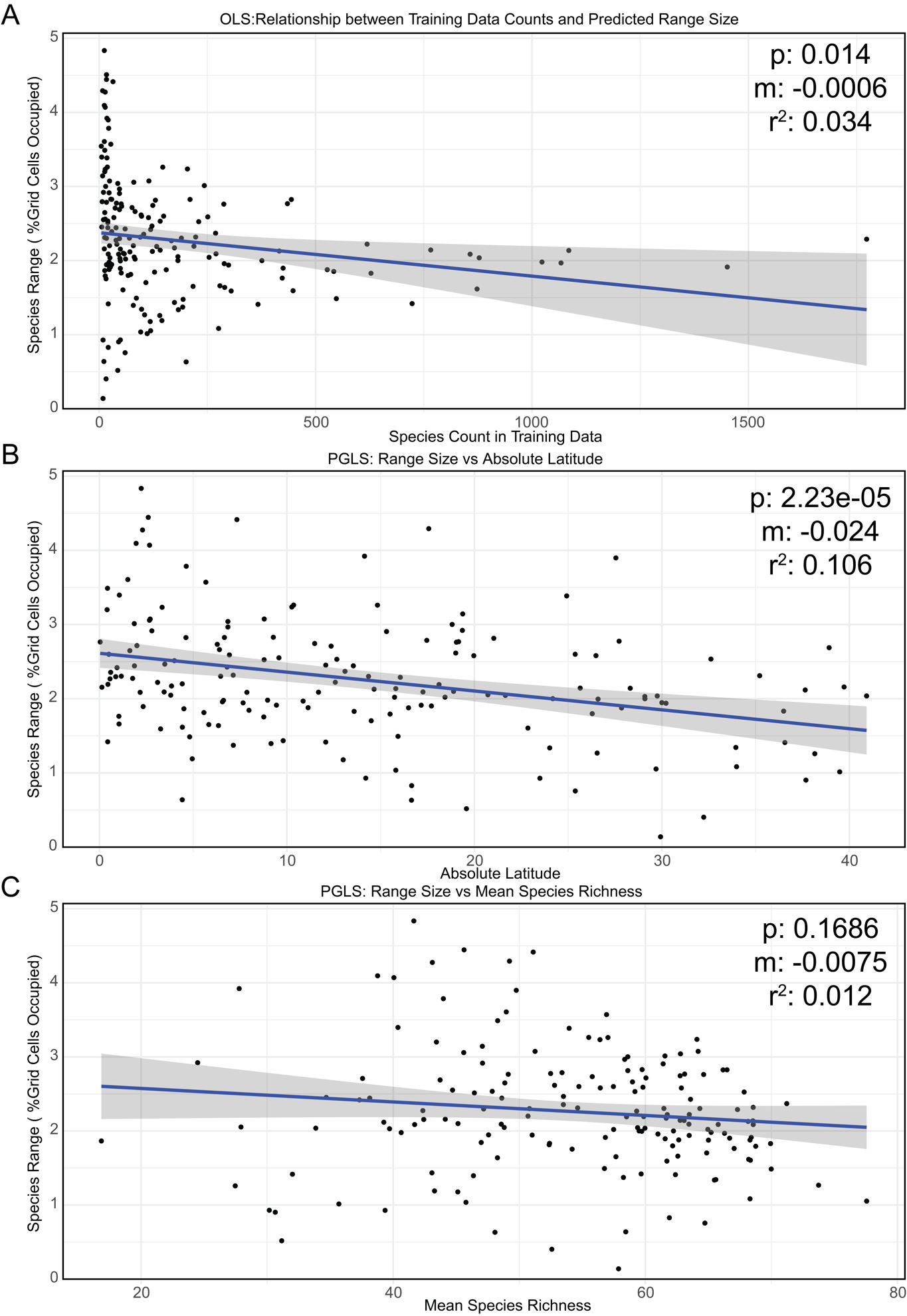

**Supplemental Figure 9:** Scatter plot of species ranges against counts in the training data, absolute latitude, and species richness. A) The X-axis is the count of occurrence records in the training, and the Y-axis is the range in percent grid cells occupied. The p-value, effect size (m), and r^2^ correspond to an Ordinary Least Squares regression of the frequency of species in the training data and their predicted range sizes. B) The X-axis is the Absolute Latitude, and the Y-axis is the range in percent grid cells occupied. The p-value, effect size (m), and r^2^ correspond to a Phylogenetic Generalized Least Squares (PGLS) analysis. C) The X-axis average species richness per species range, and the Y-axis is the range in percent grid cells occupied. The p-value, effect size (m), and r^2^ correspond to a Phylogenetic Generalized Least Squares (PGLS) analysis. Each black dot corresponds to a species, and the blue lines represent the line of fit, and the shaded region is the 95% confidence interval.

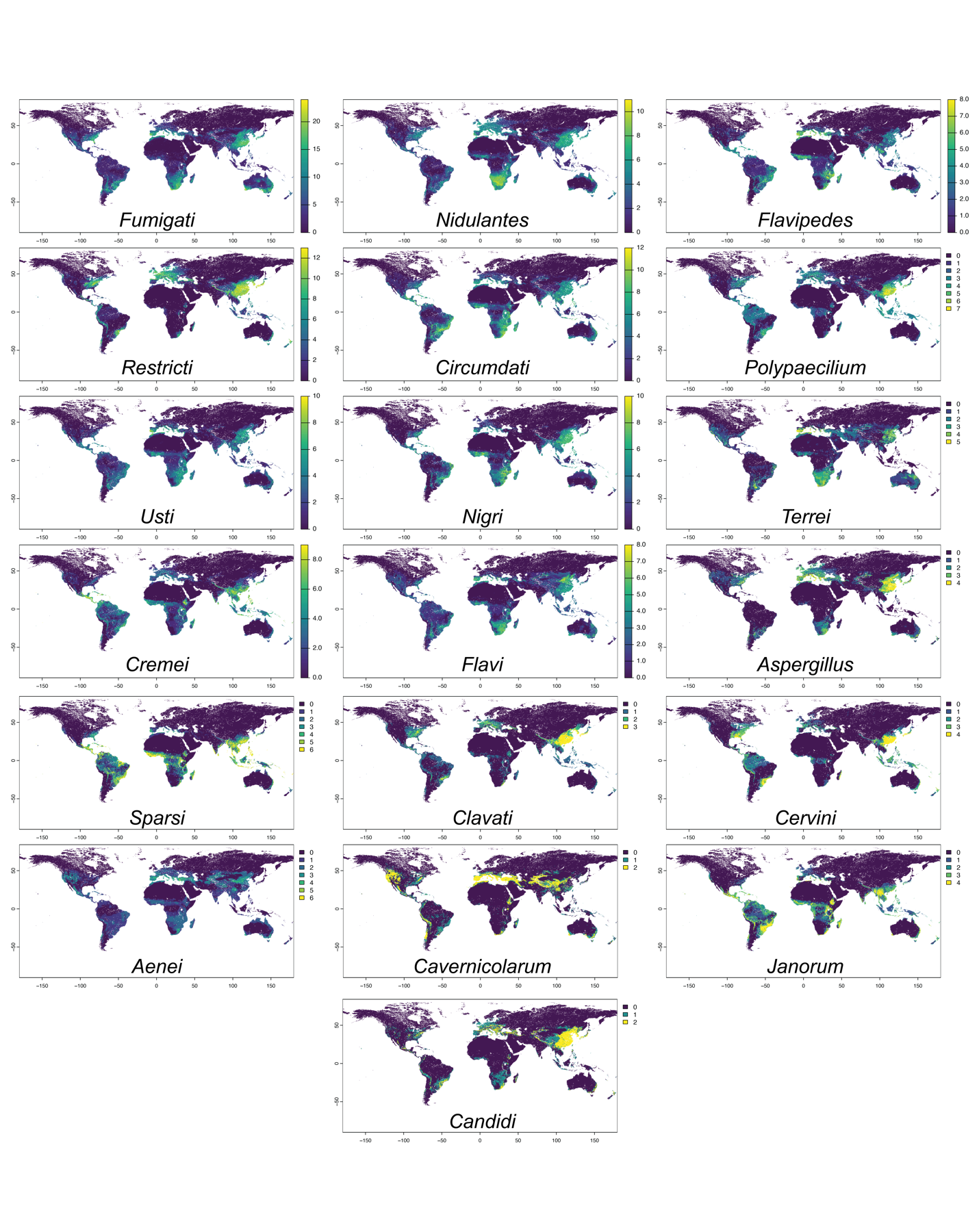

**Supplemental Figure 10:** Species richness across taxonomic sections

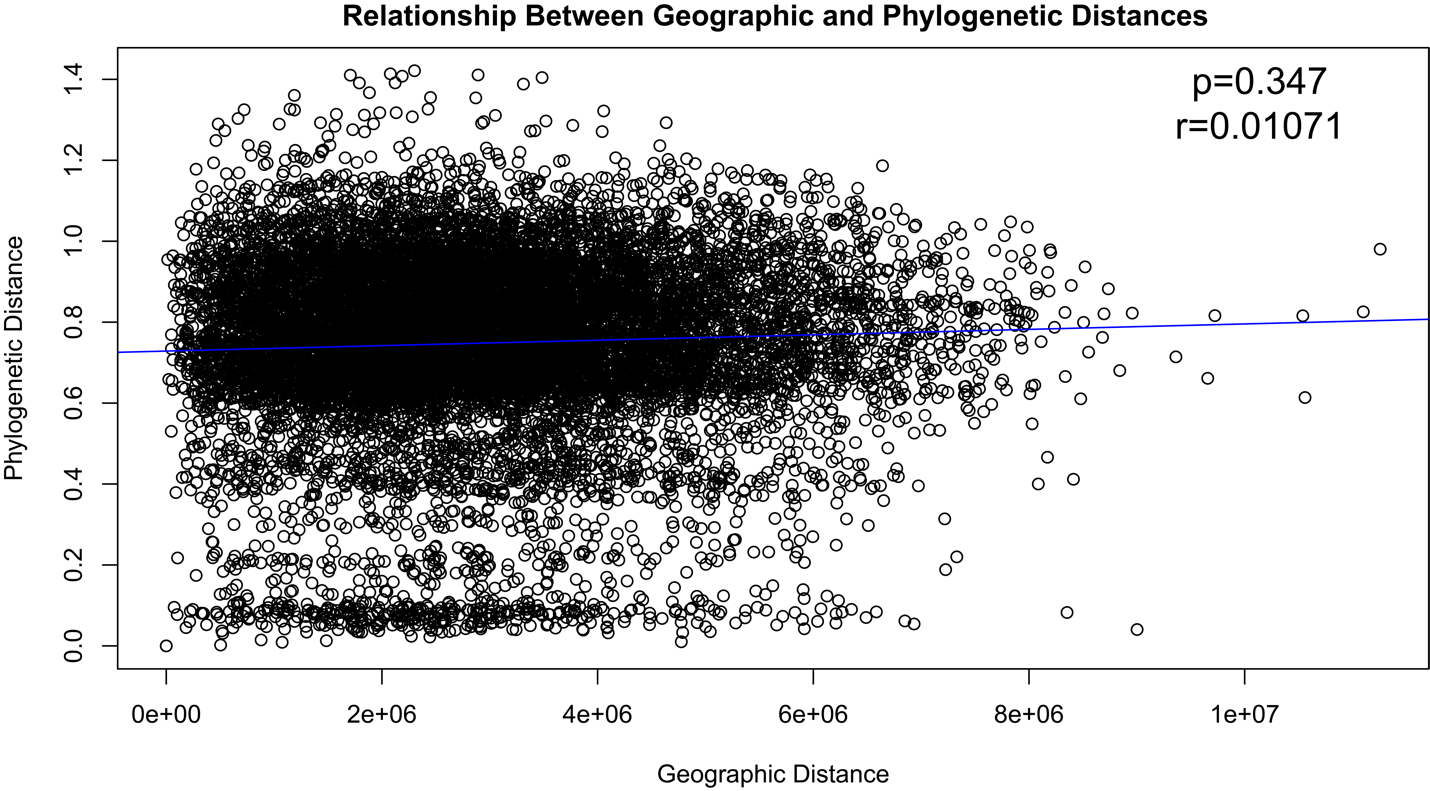

**Supplemental Figure 11:** Relationship between phylogenetic distance and geographic distance. The scatter plot depicts the correlation between geographic distance and phylogenetic distance. The X-axis represents the geographic distance in meters, and the Y-axis represents the phylogenetic distance (cophenetic distance). The blue line depicts the best-fit linear regression line. The p-value represents the statistical significance of the correlation from the Mantel test, and r is the Mantel statistic correlation coefficient.

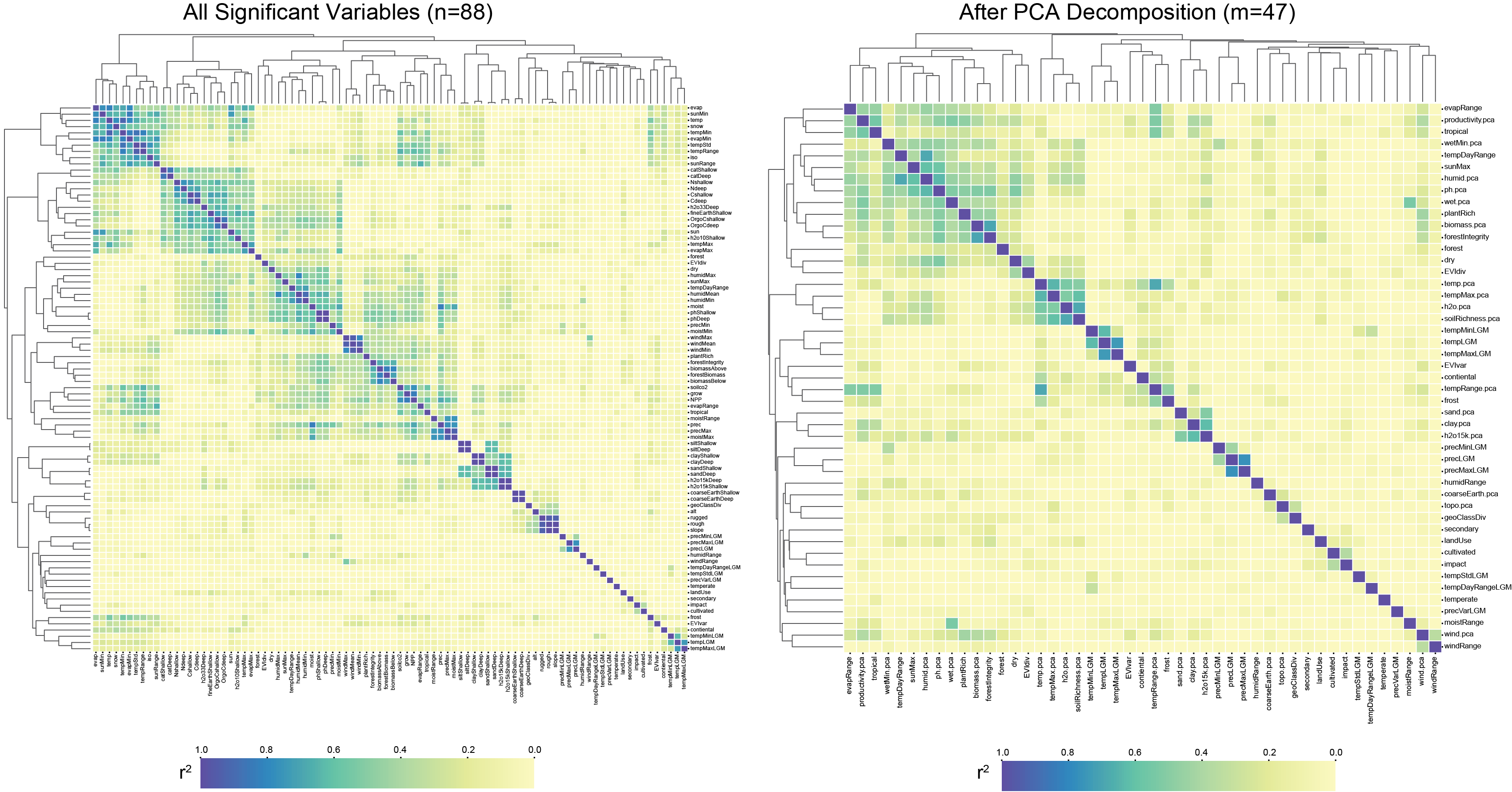

**Supplemental Figure 13:** PCA decomposition reduced the correlation between environmental variables.

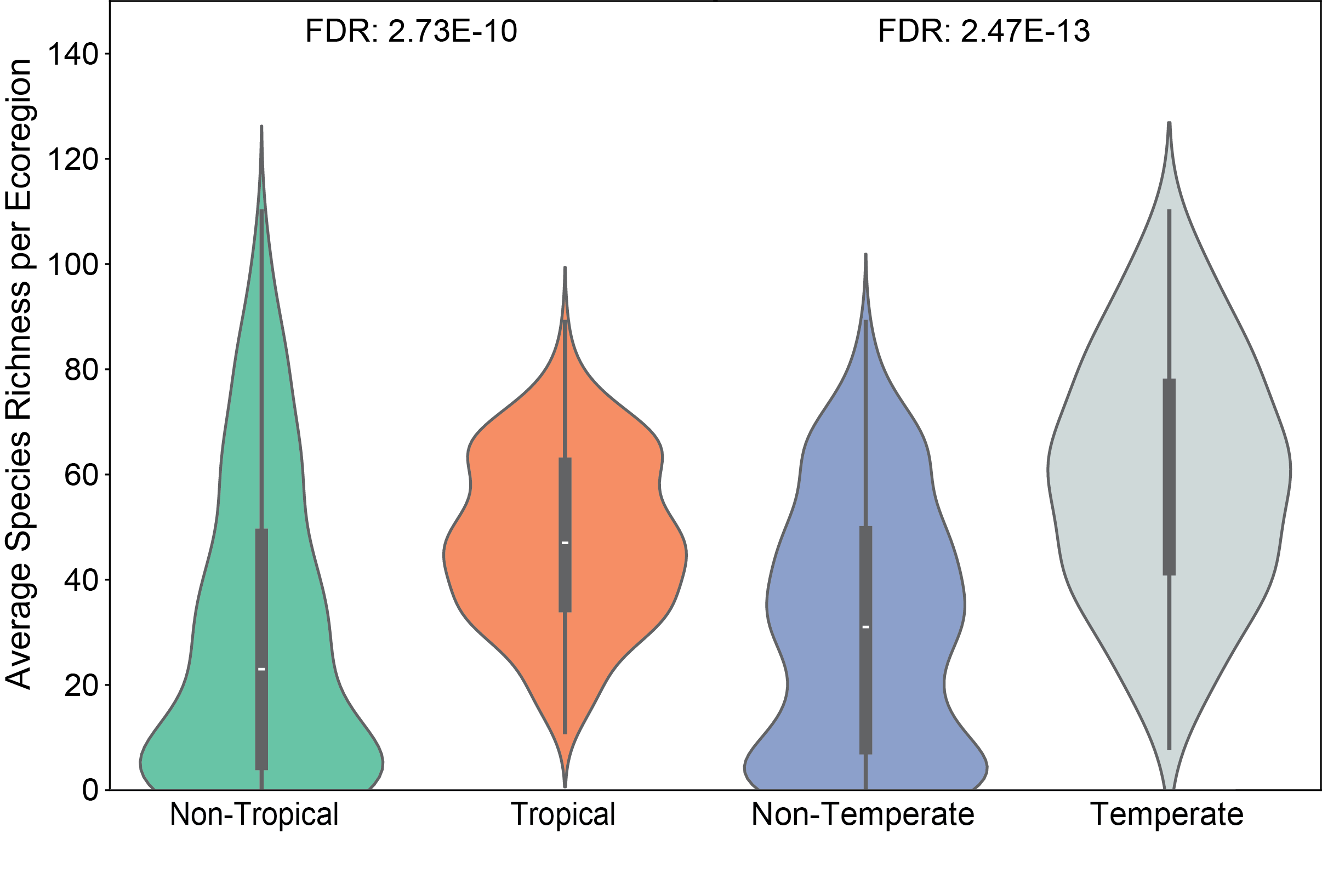

**Supplemental Figure 14:** Average species richness across ecoregions that are non-tropical, tropical, non-temperate, and temperate, and the False Discovery Rate (FDR) for the temperate and tropical categories from the negative binomial regressions.

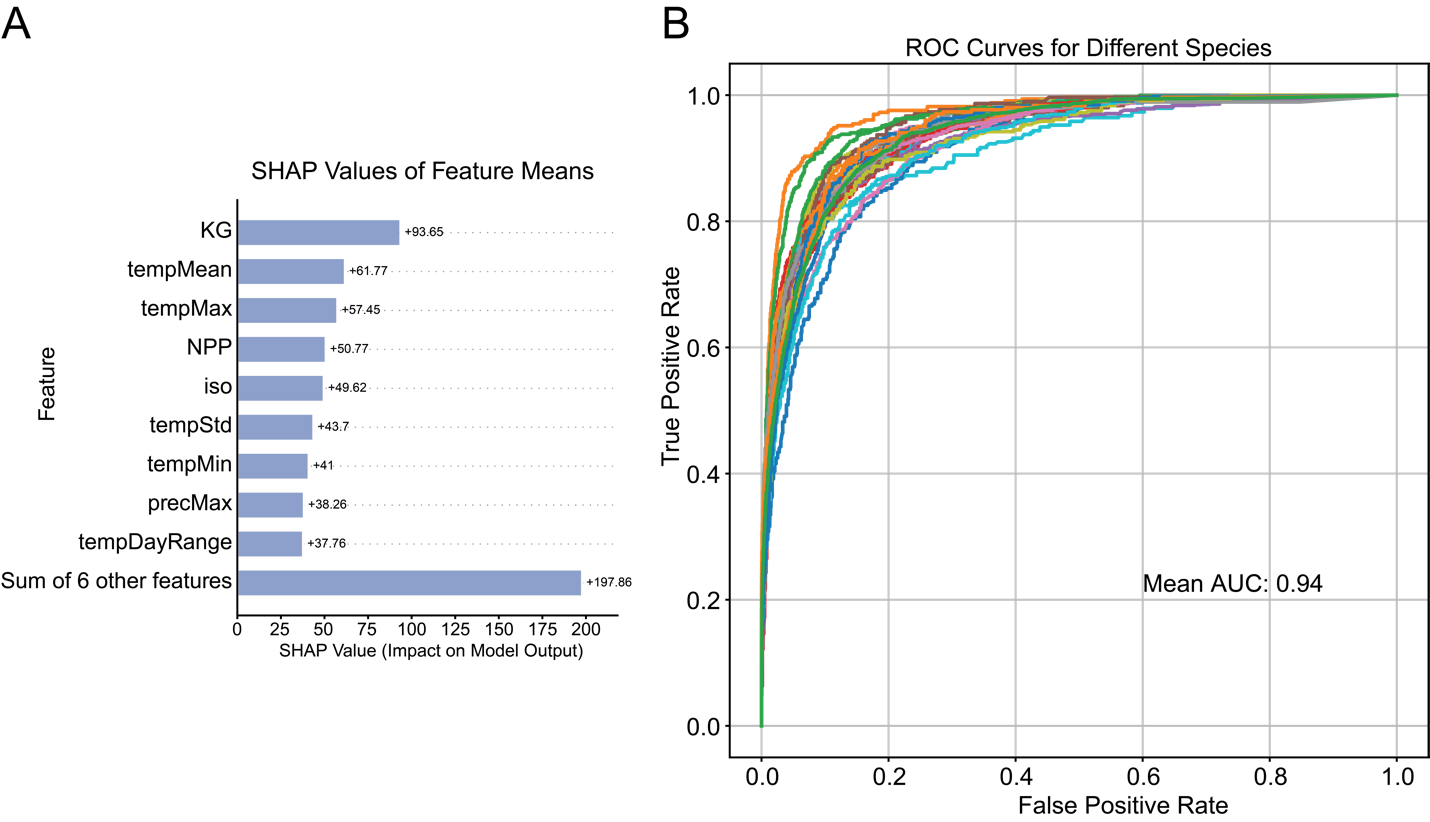

**Supplemental Figure 15:** A) Mean SHAP values for 15 features in random forest models and B) Receiver Operator Characteristic curves for 33 species with greater than 300 occurrence records.

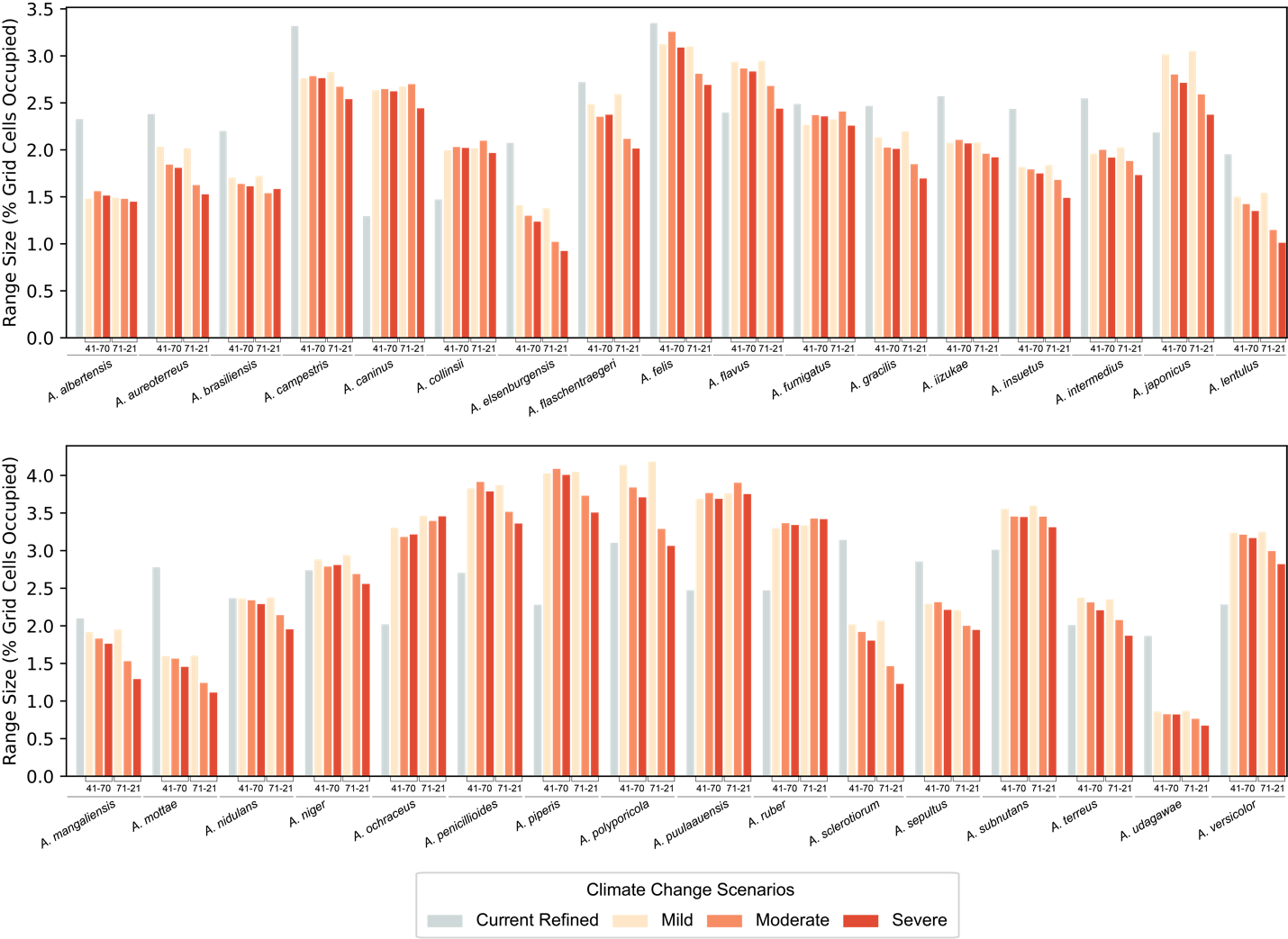

**Supplemental Figure 16:** Species predicted changes in ranges under different climate change scenarios. X-axis shows each species under the mild (sustainability, respect of environmental boundaries, and lower resource and energy intensity), moderate (regional rivalry redirecting focus to national and regional security, environmental concerns are low priority resulting in strong environmental degradation in some regions), and severe (fossil-fueled development, exploitation of fossil fuels to increase development and growth of the global economy) climate change scenarios for the two timeframes of 2041-2071 and 2071-2100. The Y-axis shows predicted ranges in percent grid cells (pixels) occupied. The grey bar is the predicted range using current data, the purple using mild scenario data, the orange bar using the moderate scenario data, and the green using severe scenario data.

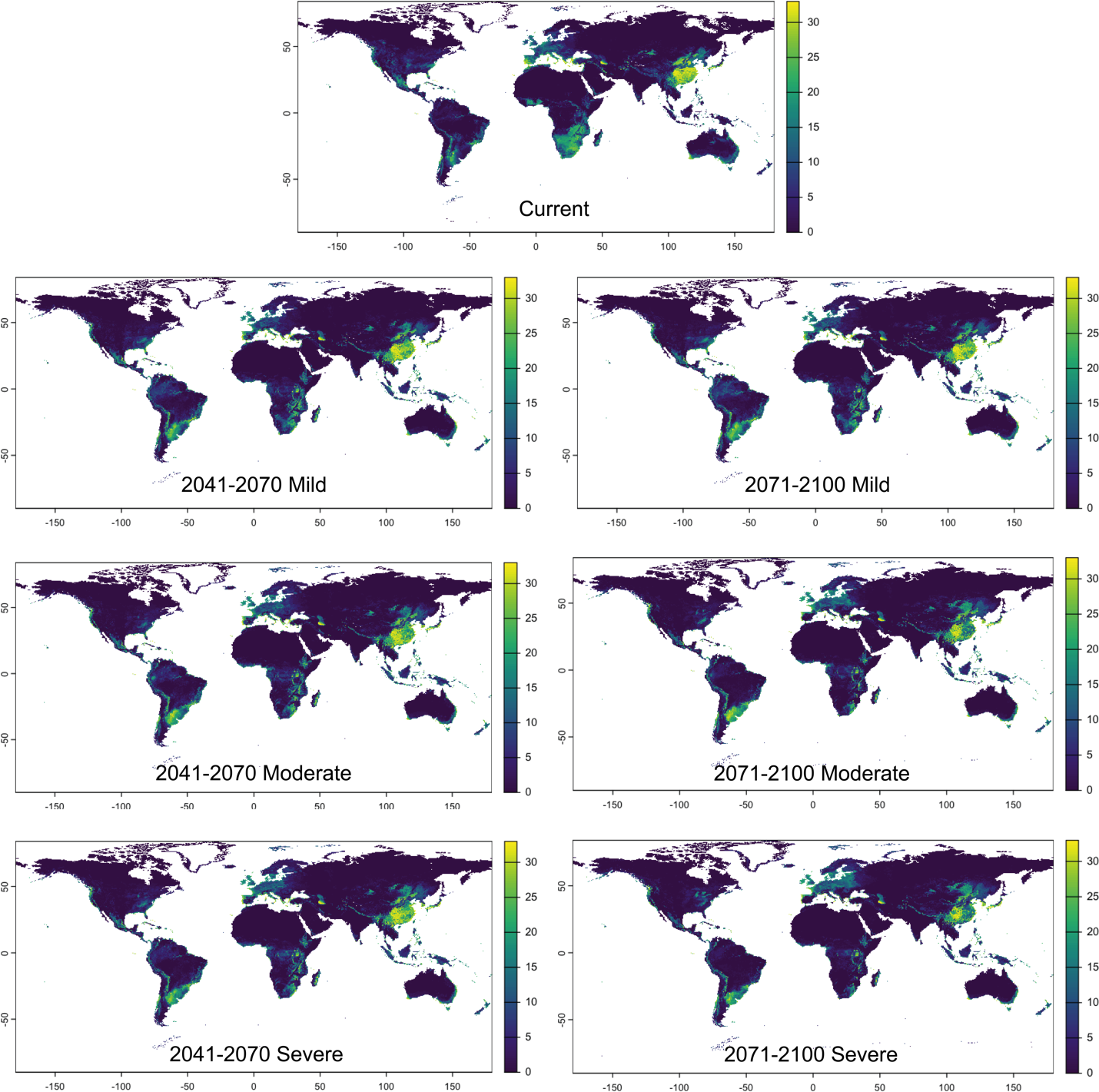

**Supplemental Figure 17:** Species richness in future predictions across both timeframes and all three climate models for 33 species.
